# Pericentrin contains internal adaptor-like sequences that activate dynein directly

**DOI:** 10.64898/2026.05.22.727342

**Authors:** Wenzhu Zhang, Ami G. Sangster, Thao T. Nguyen, John P. Gillies, Samantha Melancon, Jia-Lin Shiu, Tian Hao Huang, Akhil Gargey Iragavarapu, Xueer Jiang, Jennielee Mia, Halil Aydin, Morgan E. DeSantis, Alan M. Moses, Li-En Jao

**Affiliations:** Department of Biology, Syracuse University, Syracuse, New York, USA; Department of Cell & Systems Biology, University of Toronto, Toronto, Ontario, Canada; Department of Molecular Cellular and Developmental Biology, University of Michigan, Ann Arbor, MI, USA; Department of Molecular Pathobiology, New York University, New York, New York, USA; Department of Cell Biology and Human Anatomy, University of California, Davis, School of Medicine, Davis, California, USA; Pain Research Center, New York University, New York, NY, USA; Department of Biochemistry and Molecular Pharmacology, Grossman School of Medicine, New York University, New York, NY, USA

**Keywords:** co-translational transport, dynein, centrosome, bulk molecular features, coiled-coils, AlphaFold

## Abstract

Centrosome maturation, the rapid growth of the centrosome before cell division, prepares the cell for spindle assembly. Pericentrin (PCNT), a coiled-coil centrosomal protein, is transported to the centrosome by dynein during its translation, and this co-translational transport promotes maturation. How dynein engages PCNT remains unclear. We show that PCNT residues 1393–1525 are required for the dynein-dynactin interaction. This region shares bulk sequence features with canonical dynein cargo adaptors, and three of five synthetic sequences matching these features fully or partially restored transport of a PCNT variant lacking residues 1393–1525. PCNT is predicted to contact dynein heavy chain at sites used by canonical adaptors, and mutating these sites abolished transport. Purified PCNT activated dynein-dynactin directly without additional factors. These results suggest that PCNT binds and activates dynein through an adaptor-like sequence, with no separate adaptor, and that dynein recognizes cargoes by more diverse mechanisms than previously recognized, including an internal rather than N-terminal binding region.

## Introduction

Centrosomes function as the major microtubule organizing centers (MTOCs) of animal cells. By nucleating and organizing microtubule arrays, they help regulate cell polarity, cell signaling, and intracellular transport (Conduit et al., 2015; Barlan and Gelfand, 2017). Centrosomes also facilitate the formation of the bipolar spindle to ensure faithful chromosome segregation during mitosis (Bettencourt-Dias and Glover, 2007; Bornens, 2012). Centrosome dysfunction is linked to many human diseases, including cancers, brain disorders, dwarfism, and ciliopathies (Jaiswal and Singh, 2021).

The centrosome comprises a pair of centrioles surrounded by a proteinaceous network of pericentriolar material (PCM) (Rieder and Borisy, 1982; Vorobjev and Chentsov Yu, 1982; Wang et al., 2011; Woodruff et al., 2014; Conduit et al., 2015). The PCM contains hundreds of proteins (Andersen et al., 2003; Alves-Cruzeiro et al., 2014) and confers the MTOC activity on the centrosome (Gould and Borisy, 1977), in part because it harbors the γ-tubulin ring complexes (γ-TuRCs) that nucleate microtubules (Moritz et al., 1995; Zheng et al., 1995; Jeng and Stearns, 1999; Oegema et al., 1999). In interphase, the PCM is small, and many of its proteins are organized as nanometer-sized toroids around the mother centriole (Fu and Glover, 2012; Lawo et al., 2012; Mennella et al., 2012; Sonnen et al., 2012). As the cell enters mitosis, however, the PCM expands into a micron-sized ensemble with a concomitant increase in microtubule nucleation activity, a process termed centrosome maturation (Khodjakov and Rieder, 1999; Mahen and Venkitaraman, 2012; Piehl et al., 2004; Palazzo et al., 2000; Mennella et al., 2014).

Pericentrin (PCNT) is a large coiled-coil protein (3,336 amino acids in humans) first identified as a factor important for proper spindle organization (Doxsey et al., 1994). In vertebrates, PCNT initiates the assembly of the PCM and recruits other PCM proteins during centrosome maturation (Zimmerman et al., 2004; Haren et al., 2009; Lee and Rhee, 2011; Lawo et al., 2012). It also organizes interphase PCM. This function appears conserved among humans, mice, and flies, because loss of PCNT or its orthologs prevents the recruitment of the same set of downstream proteins in each organism (e.g., CEP215 in human, Cep215 in mice, and Cnn in flies) (Lee and Rhee, 2011; Chen et al., 2014; Lerit et al., 2015). PCNT levels are also tightly regulated. Loss-of-function mutations in *PCNT* cause microcephalic osteodysplastic primordial dwarfism type II (Rauch et al., 2008; Griffith et al., 2008; Delaval and Doxsey, 2010), whereas elevated PCNT in trisomy 21 disrupts ciliary protein trafficking and sonic hedgehog signaling, and may contribute to clinical features of Down syndrome (*PCNT* is located on chromosome 21) (Galati et al., 2018; Jewett et al., 2023).

We previously showed that human PCNT is recruited to centrosomes co-translationally during centrosome maturation (Sepulveda et al., 2018). By coupling translation to transport, this mechanism promotes the timely production and incorporation of PCNT at mitotic centrosomes (Sepulveda et al., 2018). The middle segment of PCNT (residues 854–1960) also undergoes phase separation to form dynamic condensates that move toward the centrosome in interphase cells (Jiang et al., 2021). Both the co-translational transport of endogenous PCNT and the movement of PCNT (854–1960) condensates toward the centrosome depend on the dynein motor, specifically cytoplasmic dynein 1 (Sepulveda et al., 2018; Jiang et al., 2021). How dynein engages PCNT, however, remains unclear.

Dynein is a 1.4-MDa complex assembled from two copies each of six polypeptides: the heavy chain (DHC), intermediate chain (DIC), light intermediate chain (DLIC), and three light chains (DLCs) (Trokter et al., 2012; Schmidt and Carter, 2016; Reck-Peterson et al., 2018; Canty et al., 2021). Mammalian dynein is autoinhibited and does not move processively on its own (Trokter et al., 2012; Torisawa et al., 2014; Zhang et al., 2017). Its activation requires the general cofactor dynactin (a 23-subunit, 1.2-MDa complex) and a cargo-specific adaptor (McKenney et al., 2014; Schlager et al., 2014; Carter et al., 2016; Reck-Peterson et al., 2018). Only after this tripartite dynein-dynactin-adaptor complex forms can the motor move processively along a microtubule with its cargo (McKenney et al., 2014; Schlager et al., 2014).

Approximately 20 dynein cargo adaptors have been identified to date. Despite limited sequence conservation (Reck-Peterson et al., 2018), they share two features that are critical for dynein-dynactin interactions and motor activation. The first is an N-terminal DLIC-binding (DLIC-B) domain, which folds differently among adaptor subfamilies (e.g., the CC1-Box, EF-hand, HOOK, or RH family) (Reck-Peterson et al., 2018; Olenick and Holzbaur, 2019; Singh et al., 2024). All DLIC-B domains nonetheless form a hydrophobic cleft that binds the same short C-terminal helix of DLIC (Helix-1) (Lee et al., 2018, 2020). The second is a long coiled-coil that follows the DLIC-B domain and contains two general motifs: the HBS1 motif, which binds the N-terminal tail of the DHC, and the Spindly motif, which interacts with the pointed-end subcomplex of dynactin (Urnavicius et al., 2015, 2018; Grotjahn et al., 2018; Chaaban and Carter, 2022; Singh et al., 2024). The HBS1 motif consists of a weakly conserved glutamine followed by a patch of acidic residues, which contact the conserved Y^827^ and R^759^ residues of the DHC, respectively (Chaaban and Carter, 2022; Singh et al., 2024; Aslan et al., 2026). The Spindly motif has a consensus sequence JxxEΦ in many, but not all, adaptors (J = leucine or isoleucine; Φ = small hydrophobic residue) (d’Amico et al., 2026) and binds p25 of the pointed-end subcomplex (Gama et al., 2017; d’Amico et al., 2022; Chaaban and Carter, 2022; Singh et al., 2024; Aslan et al., 2026). Because the DLIC-B domains differ among subfamilies, the HBS1 motif lacks a clear consensus, and the Spindly motif is short, new adaptors are difficult to identify by sequence homology.

Early studies in cultured cells (Purohit et al., 1999; Tynan et al., 2000; Young et al., 2000) suggested that PCNT interacts with the middle region of DLIC (Tynan et al., 2000), which differs from the region bound by the DLIC-B domains of known adaptors. These conclusions, however, rested on co-immunoprecipitation of overexpressed PCNT and/or dynein subunits from cell lysates. Whether the PCNT-DLIC interaction is direct, and whether it drives the dynein-mediated transport of PCNT, therefore remains unresolved. It is also unknown whether PCNT requires a separate adaptor to engage dynein.

To address these questions, we have combined deletion analysis, live cell imaging, quantitative proteomics, evolutionary signature analysis (Pritišanac et al., 2026), AlphaFold structural prediction (Evans et al., 2021; Jumper et al., 2021), an inducible peroxisome transport assay (Kapitein et al., 2010), and single-molecule reconstitution with purified proteins (McKenney et al., 2014; Schlager et al., 2014; Siva et al., 2026). We find that human PCNT residues 1393–1525 are required for the interaction of PCNT with dynein-dynactin and for the movement of PCNT condensates toward the centrosome. This region shares conserved bulk sequence features with the dynein-binding coiled-coils of canonical adaptors, including enrichment in glutamate and leucine and depletion of proline. Three of five synthetic sequences designed to match these features restore transport of a PCNT variant lacking residues 1393–1525, at least partially. Within this region, two HBS1-like motifs are predicted to contact the conserved Y^827^ and R^759^ residues of the DHC. In cells, PCNT (1393–1780) and PCNT (1393–1910), but not PCNT (1393–1525) alone, transport peroxisomes as efficiently as the canonical adaptor fragment BICD2N. Reversing the charge of the acidic residues in either HBS1-like motif abolishes transport. In single-molecule assays with purified dynein, dynactin, and LIS1, PCNT (1393–1780) and PCNT (1393–1910) activate processive motility with velocity and processivity indistinguishable from those of BICD2, and an excess of BICD2 competes with this activity. These results suggest that PCNT carries its own adaptor-like sequences, engages and activates dynein directly during its transport to the centrosome, without a separate adaptor.

## Results

### Human PCNT residues 1393–1525 are required for mediating PCNT-dynein interactions

We previously showed that the middle segment of human PCNT, residues 854–1960 [i.e., PCNT (854–1960)], forms dynamic condensates that move toward the centrosome in a dynein-dependent manner in retinal pigment epithelial (RPE-1) cells (Jiang et al., 2021). To identify the sequences in PCNT (854–1960) that mediate this movement, we generated a series of GFP-tagged deletion variants and followed the resulting condensates by time-lapse microscopy. Condensates formed by PCNT (854–1960) lacking residues 1393–1525 [PCNT (854–1960)Δ1393–1525] did not move toward the centrosome (**Fig. 1a,b**).

**Fig 1.**
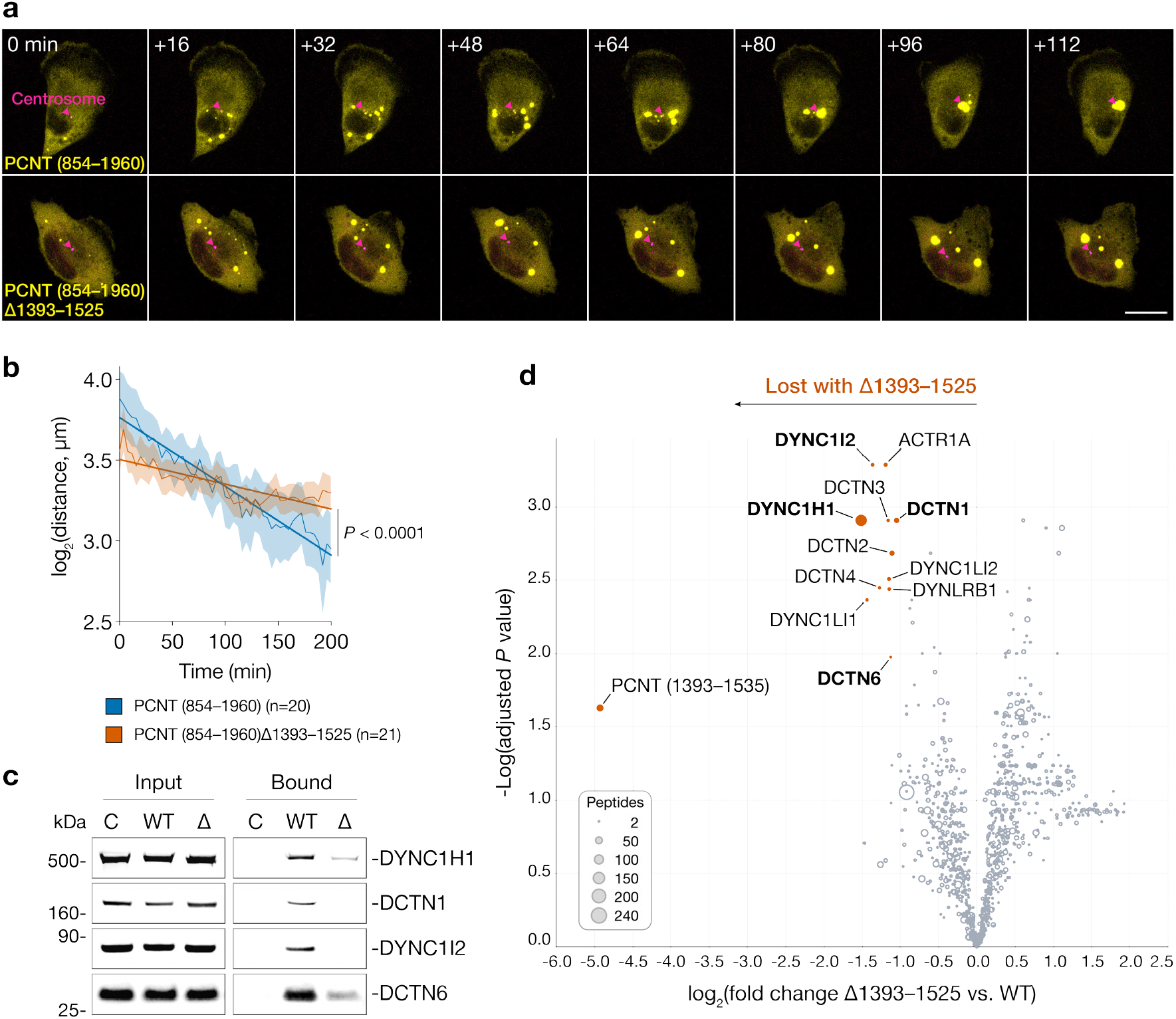
PCNT (1393–1525) is required for dynein-mediated transport of PCNT and for its interaction with dynein. **a**, Time-lapse micrographs of GFP-PCNT (854–1960) or GFP-PCNT (854–1960)Δ1393–1525 (yellow) expressed in RPE-1 cells stably expressing miRFP670nano3-CETN2 (magenta). Arrowheads denote the centrosomes. Time 0 is defined as the formation of the first condensates. Similar results were obtained in more than three biological replicates. Scale bar, 20 µm. **b**, Distance of individual condensates from the centrosome over time, aligned to the onset of phase separation (time 0) of each condensate. Data are mean ± SEM. Slopes of linear regression fits were compared by *F*-test. **c**, Anti-GFP immunoprecipitates from cells expressing GFP-FLAG (C), GFP-PCNT (854–1960)-FLAG (WT), or GFP-PCNT (854–1960)Δ1393–1525-FLAG (Δ) were analyzed by immunoblotting for dynein heavy chain (DYNC1H1), dynein intermediate chain (DYNC1I2), and the dynactin subunits DCTN1 and DCTN6. Similar results were obtained in more than three biological replicates. **d**, The immunoprecipitates in (**c**) were analyzed by quantitative 18-plex tandem mass tag (TMT) mass spectrometry from six biological replicates. Proteins de-enriched in Δ relative to WT (⩾2-fold, Benjamini–Hochberg adjusted *P* < 0.03) are shown in orange. DYNC1H1, DYNC1I2, DCTN1, and DCTN6, the proteins examined by immunoblotting in (**c**), are among them and are shown in bold. The quantified peptide is labeled PCNT (1393–1535) rather than PCNT (1393–1525) because the tryptic peptide DSQQAPLDGEVELLQQK straddles the C-terminal boundary of the deleted region (1519–1525 deleted; 1526–1535 retained) and is therefore present in the WT bait but absent from the Δ bait.

To determine whether residues 1393–1525 mediate the binding of PCNT to dynein, we first established a method for isolating native dynein-dynactin from RPE-1 cells. GFP-tagged human BICD2 (residues 25–423) (BICD2N) was used as bait in a nanobody-mediated affinity purification (Stevens et al., 2024) (**Extended Data Fig. 1**). BICD2 is among the best-characterized dynein cargo adaptors, and BICD2N has been used to isolate dynein-dynactin from cells and tissues (e.g., McKenney et al., 2014; Okada et al., 2023). Dynein heavy chain (DYNC1H1), dynein intermediate chain (DYNC1I2), and two dynactin subunits (DCTN1 and DCTN6) co-purified with BICD2N, as shown by immunoblotting (**Extended Data Fig. 1c**), validating our purification strategy of isolating dynein-dynactin complexes from RPE-1 cells.

We then purified GFP, GFP-PCNT (854–1960), or GFP-PCNT (854–1960)Δ1393–1525 by the same procedure. GFP-PCNT (854–1960) immunoprecipitates contained DYNC1H1, DYNC1I2, DCTN1, and DCTN6, but GFP and GFP-PCNT (854–1960)Δ1393–1525 immunoprecipitates contained little or none of these proteins (**Fig. 1c**).

Quantitative mass spectrometry of the same samples showed that dynein and dynactin subunits were the proteins most depleted from the GFP-PCNT (854–1960)Δ1393–1525 purification relative to GFP-PCNT (854–1960) (**Fig. 1d**). Residues 1393–1525 of PCNT are therefore required for the movement of PCNT condensates toward the centrosome and for the association of PCNT with dynein-dynactin.

### PCNT (1393–1525) and CC1 coiled-coils of dynein cargo adaptors share evolutionary signatures

Activation of mammalian dynein requires dynactin and a cargo adaptor (McKenney et al., 2014; Schlager et al., 2014; Reck-Peterson et al., 2018; Olenick and Holzbaur, 2019; Lee et al., 2020). Apart from dynein and dynactin subunits, however, our mass spectrometry analysis identified no known dynein cargo adaptor in the GFP-PCNT (854–1960) purification. This result suggests that PCNT binds dynein-dynactin through an unidentified adaptor or interacts with dynein-dynactin directly.

Cargo adaptors share limited sequence conservation (Reck-Peterson et al., 2018) yet perform the common function of activating dynein. We therefore reasoned that adaptors carry molecular features that are under selection to preserve their interaction with dynein-dynactin, and that PCNT would share these features if it binds dynein-dynactin directly. To test this hypothesis, we used a computational pipeline (Pritišanac et al., 2026, also see Methods) to calculate the “evolutionary signature” of a protein segment. An evolutionary signature is a set of z-scores, each measuring how far a bulk molecular feature of a segment (such as net charge, hydrophobicity, or the content of a particular amino acid) deviates from what is expected in the absence of selection to preserve it. A z-score far from zero indicates that selection has preserved the feature, even when the amino acid sequence itself has diverged. We computed signatures for three groups of segments. The first group comprised sixteen known dynein cargo adaptors, each divided into three segments: a B segment, which is the dynein-dynactin-binding CC1 coiled-coil between the DLIC-binding domain and the Spindly motif (Hoogenraad, 2003; Lee et al., 2020; d’Amico et al., 2022; Canty et al., 2023; Singh et al., 2024; Teixeira et al., 2025; Aslan et al., 2026), and the N and C segments, which lie N-terminal and C-terminal to the B segment, respectively. The second group comprised human PCNT, divided into three segments in the same way, with PCNT (1393–1525), the region that mediates PCNT-dynein interaction, as the B segment. The third group, which served as a negative control, comprised segments annotated as coiled-coils in UniProt (The UniProt Consortium, 2025) and supported by a publication (**Extended Data Table 1**).

PCNT (1393–1525) had an evolutionary signature similar to those of the B segments of known adaptors, but not to those of the coiled-coils outside the B segments or of most annotated coiled-coils (**Fig. 2a**). The B segments of the adaptors were distinguished by very high glutamate and leucine contents and very low proline content, and they clustered together at the upper right of the scatter plot (**Fig. 2b**). PCNT (1393–1525) also fell in this cluster, whereas most control coiled-coils did not (**Fig. 2b**). These results suggest that PCNT (1393–1525) and the CC1 coiled-coils of adaptors are under selection to preserve similar molecular features, and that PCNT (1393–1525) may interact with dynein-dynactin as cargo adaptors do.

**Fig 2.**
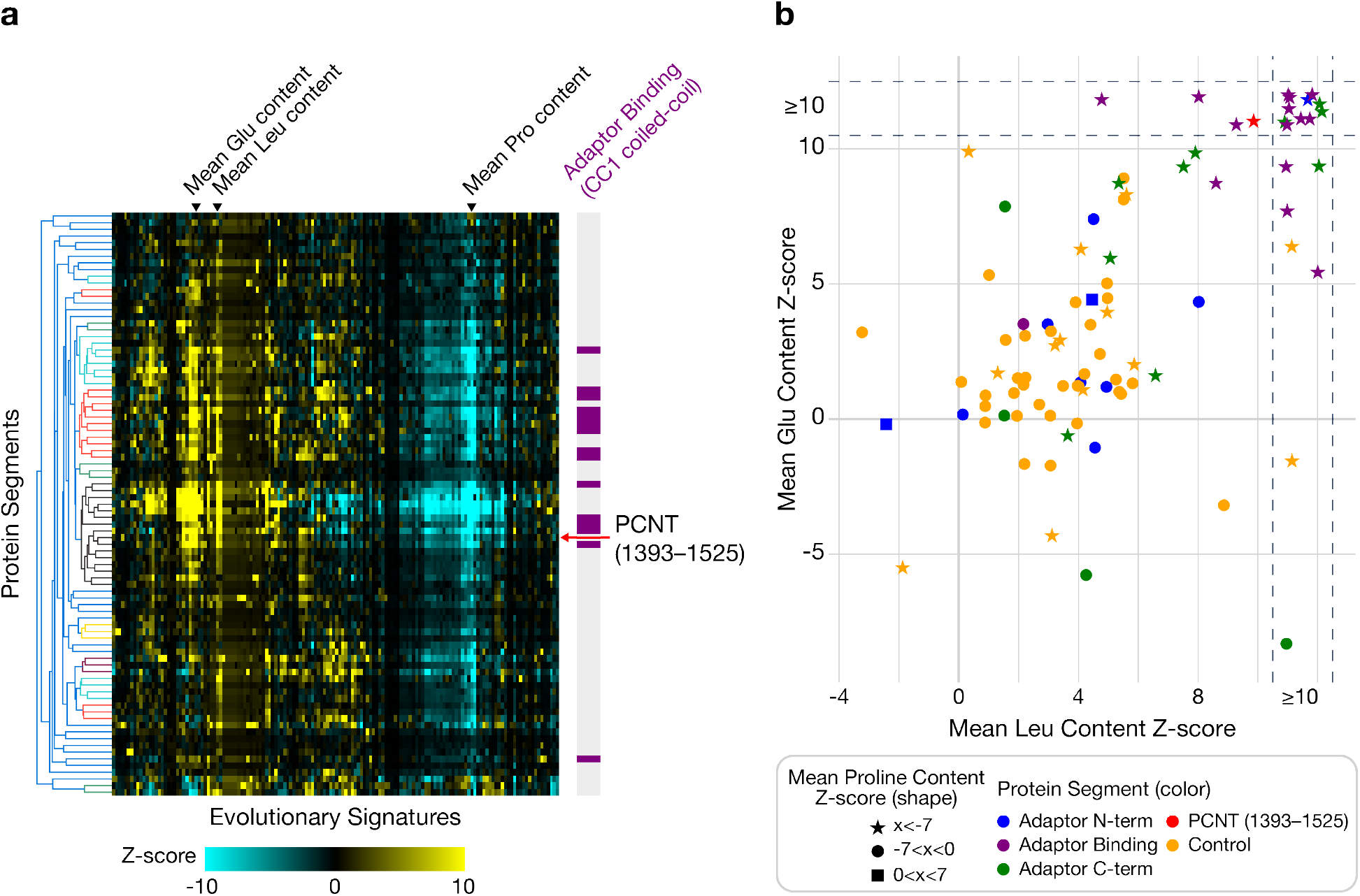
PCNT and dynein cargo adaptors share evolutionary signatures. **a**, Evolutionary signature analysis of 87 protein segments, including segments of PCNT, dynein cargo adaptors, and annotated coiled-coils from UniProt. Each row represents a segment, and each column represents the z-score of a sequence property measured in the analysis. Clusters, shown in the dendrogram to the left of the heatmap, were defined by hierarchical agglomerative clustering at a threshold of 0.4. The bar graph to the right of the heatmap marks the rows that contain the dynein-dynactin-binding segments (CC1 coiled-coils) of the cargo adaptors (purple) and the row that corresponds to PCNT (1393–1525) (red arrow). **b**, Three of the most highly conserved sequence features from the analysis, plotted for the same 87 segments. Mean leucine content z-score is shown on the x-axis, mean glutamate content z-score on the y-axis, and binned mean proline content z-score as marker shape. Colors indicate the type of segment; control refers to the annotated coiled-coils from UniProt. Z-scores were capped at ±10. Segments with z-scores ⩾10 are shown between dotted lines at the top and right of the plot, and uniform random noise was added to these points to separate them.

### Synthetic sequences matching the bulk sequence features of PCNT (1393–1525) restore dynein-mediated transport of PCNT condensates

The similarity between PCNT (1393–1525), the region required for dynein-mediated transport (**Fig. 1**), and the dynein-interacting coiled-coils of cargo adaptors (**Fig. 2**) suggested that these bulk sequence features are preserved because they allow PCNT or a cargo adaptor to engage and activate dynein. If so, designed sequences that match these features should support dynein-mediated transport even when their primary sequences differ from those of PCNT or of any adaptor. To test this prediction, we designed synthetic proteins (SPs) that match the bulk sequence features of PCNT (1393–1525) using an algorithm we developed previously (Strome et al., 2023). Starting from a coiled-coil or random sequence, the algorithm introduces point mutations iteratively and greedily retains those that bring the feature vector of the design closest to that of PCNT (1393–1525). SP1^ING4^ and SP5^STAT1^ were derived from the predicted coiled-coil regions of ING4 (residues 25–118) and STAT1 (residues 136–317), respectively. SP2, SP3, and SP4 were scaffolded from random sequences in which each amino acid was drawn with equal probability. The sequences of all five SPs are listed in **Extended Data Table 2**.

We replaced residues 1393–1525 of GFP-tagged PCNT (854–1960) with each SP, expressed the chimeras in RPE-1 cells, and assessed the movement of the resulting condensates toward the centrosome as in **Fig. 1a,b**. A chimera carrying the wild-type ING4 coiled-coil (residues 25–118), PCNT (854–1960)Δ1393–1525::ING4, did not move toward the centrosome, whereas the chimera carrying SP1^ING4^, which was derived from this ING4 sequence, did (**Fig. 3a**). Of the three designs scaffolded from random sequences, SP2 and SP3 partially restored movement (**Fig. 3b**), and SP4 did not (**Fig. 3c**). SP5^STAT1^ also failed to restore movement (**Fig. 3c**). Thus, three of the five designs restored dynein-mediated transport of PCNT condensates at least partially. These results suggest that the bulk sequence features shared by PCNT (1393–1525) and adaptor coiled-coils contribute to dynein activation, consistent with their evolutionary preservation. However, not every design that matches these features restored transport.

**Fig 3.**
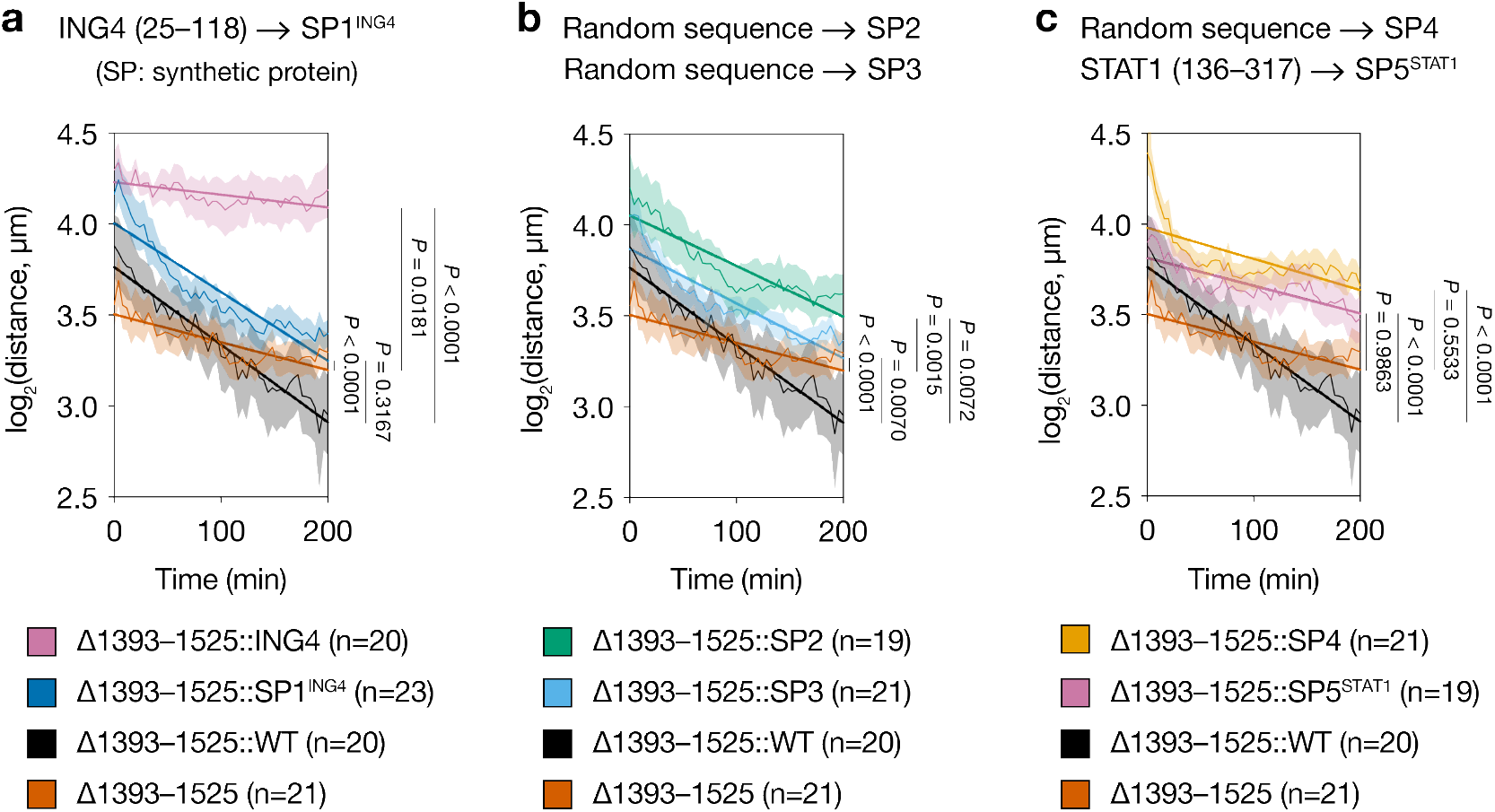
Synthetic sequences matching the bulk molecular features of PCNT (1393–1525) restore dynein-mediated transport of PCNT condensates to different extents. **a–c**, Distance of PCNT (854–1960) condensates from the centrosome over time when residues 1393–1525 were deleted (Δ1393–1525), retained (Δ1393–1525::WT), or replaced with ING4 (25–118) (Δ1393–1525::ING4) or with one of five synthetic proteins (SPs). Δ1393–1525::SP1^ING4^, Δ1393–1525::SP2, Δ1393–1525::SP3, Δ1393–1525::SP4, or Δ1393–1525::SP5^STAT1^. Each SP was designed to match the bulk molecular features of PCNT (1393–1525). Superscripts denote the protein whose predicted coiled-coil region was used to derive the SP; SP2, SP3, and SP4 were derived from random sequences. Data were collected from at least three biological replicates. Slopes of linear regression fits were compared by *F*-test, and *P* values are shown to the right of each plot. Results are summarized in **Extended Data Table 3**.

Because SP4 and SP5^STAT1^ matched the bulk features of PCNT (1393–1525) but did not restore movement, we asked what else distinguished the designs that worked. SP4 and SP5^STAT1^ had low predicted coiled-coil content (22% and 13%; **Extended Data Table 2**), and across the wild-type, deletion, and synthetic-sequence constructs, movement rate increased with predicted coiled-coil content (**Extended Data Fig. 2**). Coiled-coil content alone was not sufficient, however. ING4 (25–118) had high predicted coiled-coil content (65%) but supported little movement, whereas SP1^ING4^, which carries the bulk features of PCNT (1393–1525) and was derived from this coiled-coil scaffold (72% predicted coiled-coil content), restored movement to 89% of the wild-type rate (**Fig. 3a, Extended Data Fig. 2**, and **Extended Data Table 3**). These results suggest that restoring dynein-mediated movement requires both a high coiled-coil propensity and the bulk features shared by PCNT (1393–1525) and the adaptor CC1 coiled-coils.

### PCNT (1393–1525) contains two putative HBS1 motifs predicted to contact the dynein heavy chain

The ability of designed sequences to substitute for PCNT (1393–1525) raised the possibility that PCNT engages dynein in a manner analogous to cargo adaptors. We therefore examined PCNT (1393–1525) for the adaptor-associated motifs, specifically DLIC-binding (DLIC-B), HBS1, and Spindly motifs, and identified two HBS1-like motifs separated by approximately 40–50 amino acids (**Extended Data Fig. 3**). We refer to these as N-HBS1-like and C-HBS1-like, according to their positions in the sequence. Each motif contains a glutamine followed, approximately 15 amino acids to the C-terminus, by a cluster of acidic residues (**Extended Data Fig. 3b**). The glutamine and acidic residues are conserved in at least six vertebrates (**Extended Data Fig. 3c,d**), and this arrangement resembles that of the HBS1 motif of several adaptors (**Extended Data Fig. 3b**) (Chaaban and Carter, 2022; Aslan et al., 2026; d’Amico et al., 2026). In most cargo adaptors, cryogenic electron microscopy (cryo-EM) structures and AlphaFold2 predictions show that the glutamine and the acidic patch of the HBS1 motif contact the conserved Y^827^ and R^759^ residues of the dynein heavy chain (DHC), respectively (Sacristan et al., 2018; d’Amico et al., 2022; Chaaban and Carter, 2022; Aslan et al., 2026; d’Amico et al., 2026). We therefore used AlphaFold-Multimer (Evans et al., 2021; Jumper et al., 2021) to predict whether PCNT contacts DHC through analogous interactions. Because dynein-dynactin-adaptor complexes can contain one or two dynein dimers (Urnavicius et al., 2015, 2018), we modeled PCNT (1397–1517) with either one or two DHC (540–910) dimers. In the model with one DHC dimer, a PCNT dimer formed a complex in which Q^1466^ and the acidic patch of the C-HBS1-like motif were positioned near Y^827^ and R^759^ of the DHC, respectively (**Fig. 4b**). In the model with two DHC dimers (DHC-1 and DHC-2), the N- and C-HBS1-like motifs engaged separate DHC dimers in the same way: The glutamine residues (Q^1424^ and Q^1466^) and the acidic patches contacted Y^827^ and R^759^ of the respective DHC dimers (**Fig. 4c**). These predictions suggest that PCNT can engage dynein through its HBS1-like motifs and that its two motifs may allow it to recruit up to two dynein dimers, as reported for canonical cargo adaptors.

**Fig 4.**
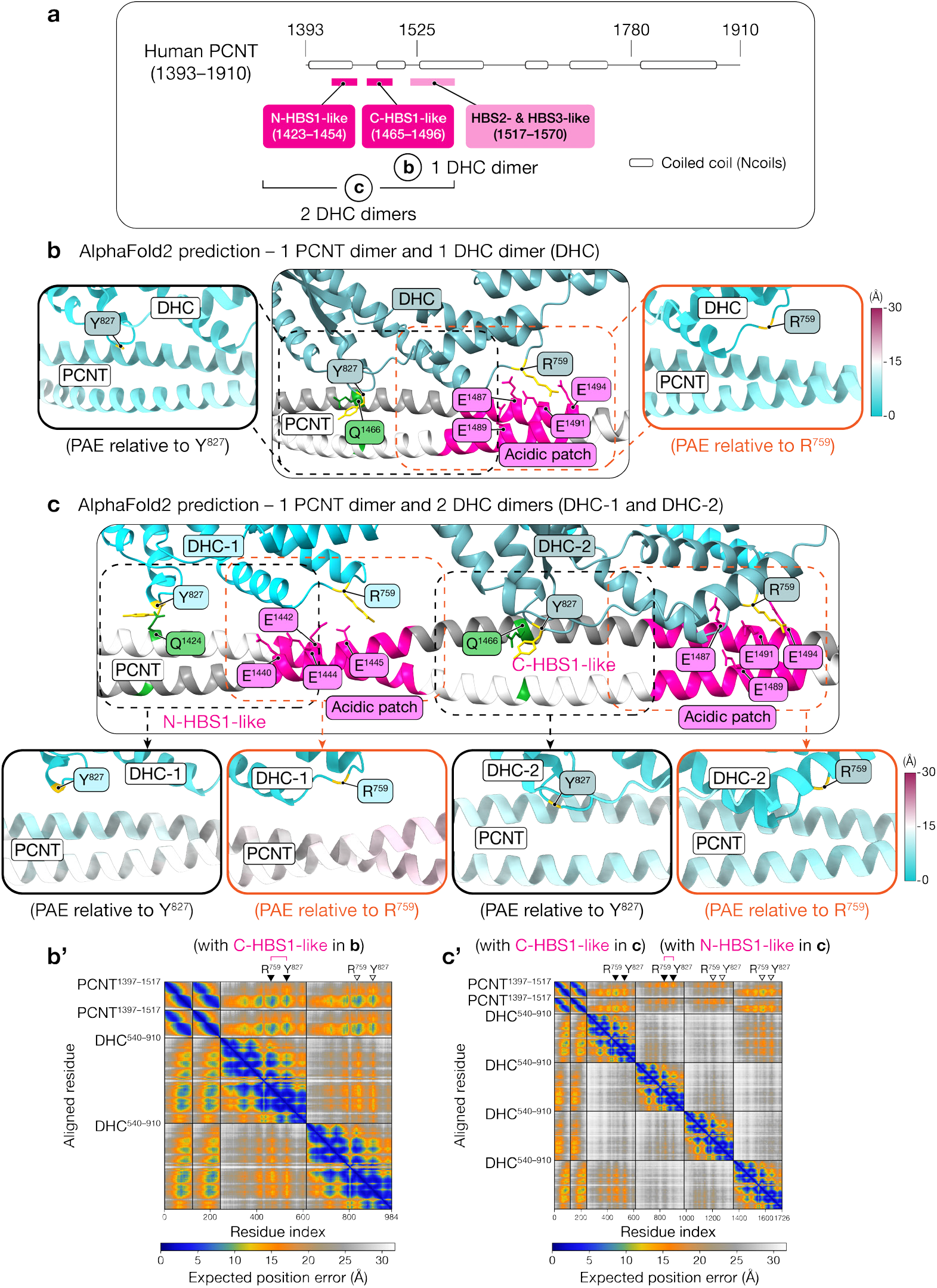
HBS1-like motifs of PCNT are predicted to interact with the dynein heavy chain. **a**, Domain structure of human PCNT (1393–1910), showing the HBS1-, HBS2-, and HBS3-like motifs (residue ranges in parentheses). Coiled-coil regions were predicted with Ncoils (Lupas et al., 1991). **b**, AlphaFold2 model of a PCNT (1397–1517) dimer with a dynein heavy chain (DHC; 540–910) dimer. Insets show the regions near Q^1466^ and the acidic patch of the C-HBS1-like motif, colored by predicted aligned error (PAE) relative to the conserved DHC residues Y^827^ and R^759^ (yellow), respectively. Lower PAE values indicate higher confidence. **c**, AlphaFold2 model of a PCNT (1397–1517) dimer with two DHC (540–910) dimers. Insets show the regions near Q^1424^ and Q^1466^ and the acidic patches of the N-HBS1- and C-HBS1-like motifs, colored by PAE relative to DHC residues Y^827^ and R^759^ (yellow), respectively. The full PAE plots are shown in (**b’**) and (**c’**); solid arrowheads mark the positions of Y^827^ and R^759^, the residues shown in the models in (**b**) and (**c**).

### PCNT acts like an activating dynein cargo adaptor in cells

The AlphaFold2 models predict contact between PCNT and dynein, but they do not show whether this contact activates dynein. To test activation, we used an inducible peroxisome transport assay in cells (Hoogenraad, 2003; Kapitein et al., 2010). In this assay, we expressed a peroxisome-targeting sequence (Kapitein et al., 2010) fused to mStayGold (mSG1) (Ando et al., 2023) and FK506 binding protein (FKBP) (Huynh and Vale, 2017; Wang et al., 2019), together with BICD2N or PCNT sequences fused to mScarlet3 (mSL3) (Gadella et al., 2023) and the FKBP12-rapamycin-binding domain (FRB) (**Fig. 5a**). Rapamycin induces heterodimerization of FKBP and FRB and thereby brings peroxisomes together with the BICD2N or PCNT fusion protein. If a construct acts as a dynein activating adaptor, the peroxisomes it recruits should move toward the centrosome.

**Fig 5.**
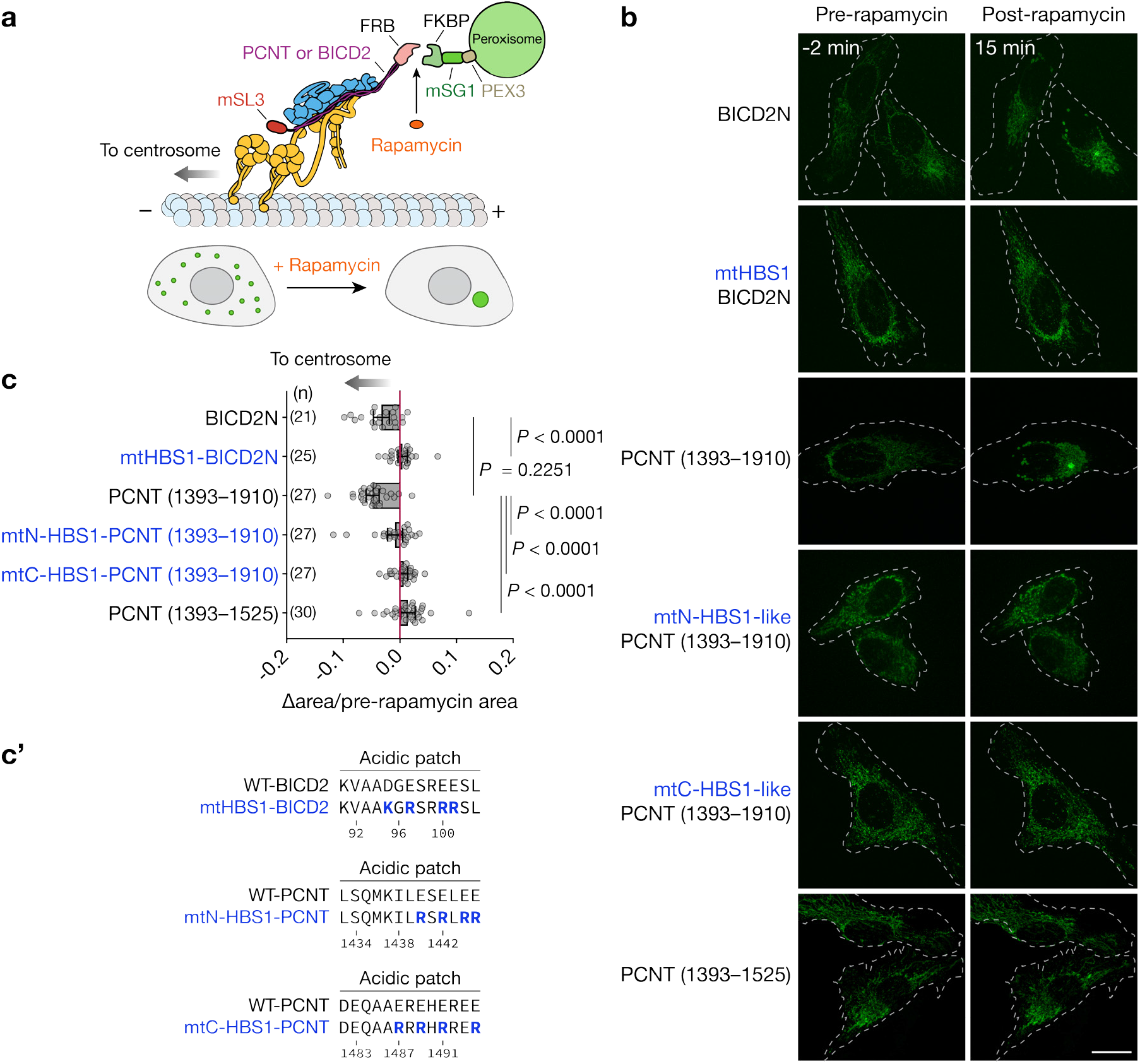
PCNT (1393–1910) acts like a canonical dynein cargo adaptor in cells. **a**, Schematic of the inducible peroxisome transport assay (adapted from Aslan et al., 2026). HeLa cells were transiently transfected with BICD2 or PCNT variants fused to mScarlet3 (mSL3)-FRB and with a peroxisome-targeting PEX3-mStayGold (mSG1)-FKBP fusion. Rapamycin induces FRB-FKBP dimerization, which localizes the dynein-dynactin-adaptor complex to peroxisomes and drives their retrograde transport toward the microtubule minus end. **b**, Representative images of the peroxisome transport assay. Cells were imaged 2 minutes before and 15 minutes after the addition of rapamycin (time 0). Scale bar, 20 µm. **c**, Quantification of peroxisome motility. Each data point is the change in mSG1-positive area per cell (15 min post-minus pre-rapamycin) divided by the pre-rapamycin mSG1-positive area. Peroxisomes were scored as trafficking if this ratio was negative and the post-rapamycin area was significantly smaller than the pre-rapamycin area (two-tailed paired *t*-test). Data are mean ± 95% CI from at least three biological replicates. *P* values were calculated by one-way ANOVA with Dunnett’s test relative to the BICD2N or PCNT (1393–1910) condition. The number of cells quantified (n) for each condition is shown in parentheses. **c’**, Locations of the charge-reversal mutations in the acidic patches of the BICD2 HBS1 and PCNT HBS1-like motifs. See **Extended Data Fig. 3b** for the full sequences of the motifs.

BICD2N, the positive control, transported peroxi-somes efficiently to the centrosome upon rapamycin addition (**Fig. 5b,c**). PCNT (1393–1525) did not induce peroxisome transport. Longer fragments, PCNT (1393–1780) (**Extended Data Fig. 4d**) and PCNT (1393–1910) (**Fig. 5b**,**c**), however, were as efficient as BICD2N. To test whether transport depends on the HBS1-like motifs predicted by the AlphaFold models (**Fig. 4c**), we reversed the charges of the acidic residues predicted to contact DHC R^759^ in either the N-HBS1-like motif (^1440^ESELEE^1445^-to-RSRLRR) or the C-HBS1-like motif (^1487^EREHEREE^1494^-to-RRRHRRER) (**Extended Data Fig. 3b**). Either substitution abolished peroxisome transport (**Fig. 5b,c**,**c’**). Reversing the charges in the HBS1 motif of BICD2 had the same effect (**Fig. 5b,c,c’**), indicating that this substitution disrupts dynein activation by a known adaptor. Together, these results indicate that PCNT can activate dynein in cells and that this activity requires the acidic patches of both HBS1-like motifs.

### PCNT activates processive motility of purified dynein-dynactin complexes

The peroxisome assay shows that PCNT can transport cargo toward the centrosome in cells, but it does not show that PCNT activates dynein directly, because other cellular factors could contribute. To test direct activation, we reconstituted dynein-dynactin-activator motility from purified components, the standard assay for dynein activators (McKenney et al., 2014; Schlager et al., 2014). We purified human cytoplasmic dynein 1 complex and dynein regulator LIS1 from insect cells and dynactin from HEK293T cells, each carrying the tags required for purification and fluorescent labeling (**Extended Data Fig. 5e**). HaloTag-tagged BICD2 (1–598), PCNT (1393–1780), and PCNT (1393–1910) were purified from *E. coli* (**Extended Data Fig. 5e**).

We assembled and visualized single dynein-dynactin-activator complexes moving on microtubules by two-color total internal reflection fluorescence (TIRF) microscopy. Without an activator, dynein-dynactin was largely stationary, with occasional diffusive events and rare processive runs (**Fig. 6a,b** and **Extended Data Fig. 5a,b**). As expected, BICD2 activated processive movement of dynein-dynactin and comigrated with it (**Fig. 6**). PCNT (1393–1780) and PCNT (1393–1910) did the same (**Fig. 6**), although the landing rate of processive events was lower for PCNT than for BICD2 (**Fig. 6b**). Once PCNT-dynein-dynactin complexes began to move, their processivity and velocity were indistinguishable from those of BICD2-dynein-dynactin complexes (**Fig. 6c,d**), and their run lengths were slightly longer (**Fig. 6e**).

**Fig 6.**
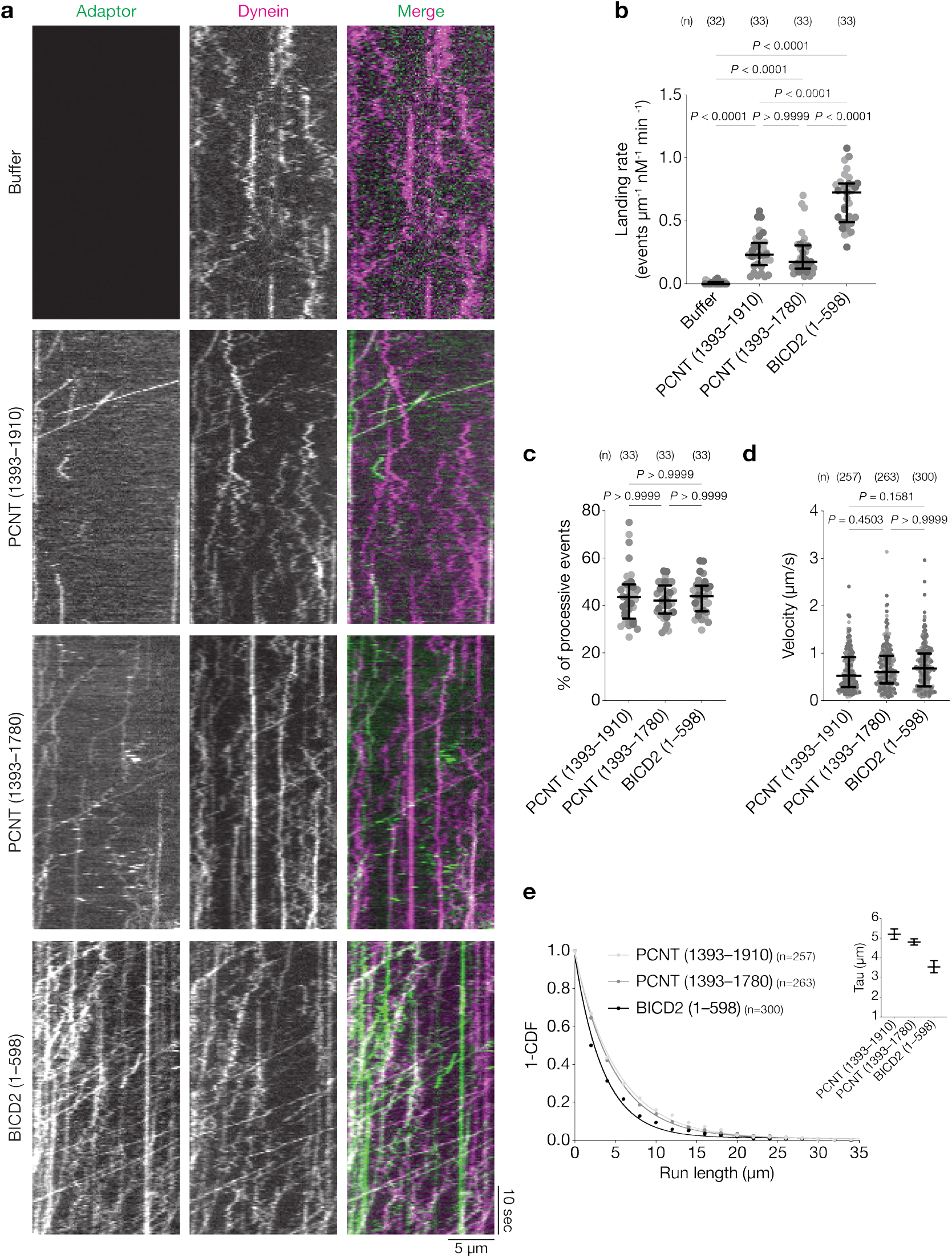
PCNT is a dynein activator. **a**, Representative kymographs of the adaptor, dynein, and merged channels for complexes containing dynein, dynactin, and LIS1 in the presence of buffer alone, PCNT (1393–1910), PCNT (1393–1780), or BICD2 (1–598). **b**, Landing rate of processive adaptor events (events per µm of microtubule per nM dynein per minute). Each spot represents one microtubule, and shades of gray distinguish the three replicates, which came from two independent preparations of adaptor and dynein. Error bars indicate median ± interquartile range. **c**, Percentage of adaptor events that were processive. Data are presented as in (**b**). **d**, Velocity of motile adaptor events (µm/s). Each spot represents one event; replicates and statistics are as in (**b**). **e**, One minus the cumulative distribution function (1-CDF) of adaptor run lengths for all motile complexes. Inset, characteristic run length (Tau) with 95% confidence intervals, obtained by fitting the 1-CDF to a one-phase decay. Statistical significance in **b**–**d** was assessed by a Kruskal-Wallis test with Dunn’s multiple comparisons test. *P* values for pairwise comparisons are shown above the plots, and the number of samples quantified (n) for each condition is shown in parentheses.

PCNT could activate dynein itself, or it could instead bind dynein-dynactin complexes that have already been activated by another factor. To distinguish these possibilities, we added an excess of unlabeled BICD2 to the reactions. This addition abolished the processive events of labeled PCNT (**Extended Data Fig. 5c,d**), indicating that PCNT and BICD2 compete for dynein, as expected if both act as activators. These results indicate that PCNT activates dynein-dynactin directly as canonical dynein activators do.

## Discussion

Centrosome maturation ensures faithful chromosome segregation and is facilitated by the dynein-mediated, co-translational transport of pericentrin (PCNT) to the centrosome in human cells. PCNT then recruits additional centrosomal proteins and greatly increases the microtubule-nucleating activity of the centrosome. Here we have defined the basis of this transport by combining cellular, biochemical, and computational approaches. We find that PCNT is itself a dynein activator. Residues 1393–1525 of PCNT are required for dynein-dynactin interaction and share bulk sequence features with the dynein- and dynactin-binding coiled-coils of known cargo adaptors, and designed sequences matching these features restored transport. Structural models predict that this segment engages the dynein heavy chains (DHCs) through sequences resembling the HBS1 motif of cargo adaptors. PCNT (1393–1525) alone did not support peroxisome transport, but the longer segments PCNT (1393–1780) and PCNT (1393–1910), which contain additional coiled-coils, drove transport as efficiently as BICD2N, the N-terminal fragment of the canonical adaptor BICD2. Both segments also formed processive complexes with purified dynein and dynactin in vitro. These results indicate that PCNT carries its own adaptor-like sequences, which engage and activate dynein-dynactin without a separate adaptor.

### PCNT (1393–1525) and the CC1 coiled-coil of dynein cargo adaptors are under similar selection pressure

PCNT (1393–1525) shares evolutionary signatures with the CC1 coiled-coils of sixteen known dynein cargo adaptors. These regions have conserved bulk sequence features: they are enriched in charged and acidic residues, especially glutamate, and in leucine, and depleted of glycine, polar residues, and proline (**Fig. 2**). In contrast, the regions N- or C-terminal to the adaptor CC1 coiled-coils have different evolutionary signatures. Both PCNT (1393–1525) and the adaptor CC1 coiled-coils also have a high propensity to form coiled-coils. However, this propensity cannot be the only feature under selection, because most coiled-coil sequences lack these bulk sequence features (**Fig. 2**). Instead, our data suggest that these sequences are selected for their ability to engage and activate the dynein-dynactin complex. Consistent with this model, de novo synthetic sequences designed from these conserved features at least partially restored dynein-mediated transport of PCNT condensates if they also have a high propensity to form a coiled-coil (**Fig. 3** and **Extended Data Table 3**). Thus, supporting dynein movement requires coiled-coil-forming propensity and the bulk sequence features of PCNT (1393–1525), and both properties are maintained despite extensive changes in primary amino acid sequence (**Fig. 3** and **Extended Data Table 3**). This conservation of bulk molecular features, rather than primary sequence, has two implications. First, the binding interface formed by dynein and dynactin may have a high degree of conformational flexibility, allowing it to accommodate diverse adaptors and, possibly, certain cargoes directly (e.g., PCNT). Second, converting ING4 into SP1^ING4^ changed the function of the coiled-coil without markedly changing its predicted coiled-coil content (65% and 72%): ING4 supported little movement, whereas SP1^ING4^ restored nearly wild-type motility. Despite their simple architecture, coiled-coils are among the most versatile and diverse protein folds in nature (Lupas et al., 2017; Burkhard et al., 2001). Bulk sequence features may therefore be an underappreciated way to tune their functions, well beyond dynein activation.

### PCNT binds one or two DHC dimers through two HBS1-like motifs

In our models, PCNT engages DHC through two HBS1-like motifs (**Fig. 4**), whereas most cargo adaptors have a single HBS1 motif. As in other cargo adaptors (Urnavicius et al., 2015; Chaaban and Carter, 2022), a PCNT dimer can engage with one or two DHC dimers. With one DHC dimer, the C-HBS1-like motif is predicted to mediate the interaction. With two, the two motifs are predicted to engage separate DHC dimers. In both cases, a glutamine and the acidic patch of the HBS1-like motif contact the conserved Y^827^ and R^759^ residues of DHC, respectively, an arrangement that resembles the HBS1-DHC interface of canonical cargo adaptors (Sacristan et al., 2018; Chaaban and Carter, 2022; d’Amico et al., 2022; Canty et al., 2023). Reversing the charges of four glutamates around R^759^ in either motif abolished PCNT-mediated peroxisome transport (**Fig. 5**). This result suggests that the transport depends on the predicted PCNT-DHC interface and not on an indirect mechanism, and that the interface relies on electrostatic interactions.

While this manuscript was in preparation, d’Amico et al. reported cryo-EM structures in which the cargo adaptor HOOK3 engages two DHC dimers through two consecutive HBS1 motifs (d’Amico et al., 2026). Among the adaptors they examined, only HOOK3 has two HBS1 motifs. These motifs are separated by approximately 40–50 amino acids, a spacing similar to that of the two HBS1-like motifs of PCNT (**Extended Data Fig. 3**). d’Amico et al. also found that most adaptors contact DHC through additional charged interactions downstream of HBS1, which they termed HBS2 and HBS3, spaced approximately 30 amino acids apart (d’Amico et al., 2026). PCNT contains two acidic patches, conserved in six vertebrates, located approximately 30–60 amino acids C-terminal to the C-HBS1-like motif (**Extended Data Fig. 3d**). PCNT and HOOK3 may therefore share a similar arrangement of HBS1, HBS2, and HBS3 motifs. Whether PCNT engages DHC through the same mechanism as HOOK3 will become clearer when structures of PCNT-DHC complexes are determined.

### The dynein-dynactin binding sites after PCNT (1393–1525)

Although PCNT (1393–1525) contains the HBS1-like motifs that engage DHC, it did not drive peroxisome transport on its own. PCNT (1393–1780) (**Extended Data Fig. 4d**) and PCNT (1393–1910) (**Fig. 5**) did so efficiently, so the additional sequence beyond residue 1525 must contribute. In other adaptors, a Spindly motif supports a stable adaptor-dynein-dynactin complex by contacting the pointed end of dynactin, and PCNT may contain a similar motif in this region. To test this, we substituted alanine for the critical leucine and glutamate residues at positions 1, 4, and 5 (3A mutations) in several candidate motifs. The same mutation in the Spindly motif of BICD2N abolished peroxisome transport (**Extended Data Fig. 4d,d’**). The best candidate in PCNT (residues 1771–1775) lies near a break between predicted coiled-coils, a feature of many Spindly motifs (d’Amico et al., 2022; Chaaban and Carter, 2022; d’Amico et al., 2026). Nevertheless, the 3A mutation at this site did not disrupt peroxisome transport, and none of the other candidates did either (**Extended Data Fig. 4** and data not shown).

Two explanations are possible. First, we may not have mutated the correct motif. Atypical Spindly-like motifs that diverge from the consensus have been reported (Aslan et al., 2026; d’Amico et al., 2026), and PCNT may contain one that cannot be recognized from sequence alone. Second, we may have mutated the correct motif, but it may be dispensable in the peroxisome assay. Contact with the pointed end of dynactin is not required for all adaptors; for example, deleting the Spindly motif of JIP3 has minimal effects on dynein motility in vitro (Singh et al., 2024). In addition to Site 4, where p25 contacts the Spindly motif, the pointed end of dynactin offers additional adaptor-binding sites (e.g., Site 2 and Site 3, on p62 and p25) (Lau et al., 2021). If adaptors differ in how much they depend on these additional sites, disrupting the Spindly motif alone could impair motility mediated by some adaptors (for example, BICD2; **Extended Data Fig. 4**) but not others.

The HOOK3 structures suggest a further reason why the Spindly motif of PCNT may be dispensable. d’Amico et al. found that two HOOK3 dimers engage the dynein-dynactin complex (d’Amico et al., 2026). If PCNT likewise uses two dimers, in addition to two HBS1-like motifs, it may contact the pointed end of dynactin at multiple sites, and disrupting the Spindly motif alone would then not abolish the PCNT-dynein-dynactin interaction. Testing this model will require mutations at other potential contact sites around the pointed end, and structures of PCNT bound to the pointed-end complex.

### The dynein-dynactin binding sites before PCNT (1393–1525)

No known DLIC-B domain (e.g., CC1-Box, HOOK, EF-hands) is evident N-terminal to the HBS1-like motifs of PCNT (1393–1910). Nevertheless, PCNT (1393–1780) and PCNT (1393–1910) move peroxisomes toward the centrosome as efficiently as BICD2N, which contains the DLIC-B domain CC1-Box. In the single-molecule assay, complexes of dynein and dynactin with either PCNT fragment also showed processivity and velocity indistinguishable from those of BICD2 complexes (**Fig. 6c,d**). Two explanations are possible. PCNT may contain an atypical DLIC-B domain upstream of the HBS1-like motifs, or it may activate dynein-dynactin without binding DLIC. The second is plausible because the adaptors NINL and BICD2 can still activate dynein motility in vitro without their DLIC-B domains, although less efficiently (Siva et al., 2026). Two experiments would distinguish these possibilities. The first tests whether PCNT binds DLIC directly. The second tests whether motility of PCNT-dynein-dynactin complexes persists when the DLIC-B interaction is blocked, either by adding excess DLIC Helix-1, the C-terminal helix that binds the DLIC-B domain of adaptors, or by mutating its two key phenylalanine residues (Lee et al., 2018). Persistent motility would support activation without DLIC binding.

### Why do PCNT and some adaptors engage with dynein co-translationally?

Dynein moves PCNT toward the centrosome cotranslationally (Sepulveda et al., 2018; Safieddine et al., 2021). Three other bona fide dynein cargo adaptors (NUMA1, NIN, and BICD2) and one putative adaptor (CCDC88C) (Redwine et al., 2017) are also transported to the centrosome co-translationally (Chouaib et al., 2020; Safieddine et al., 2021), presumably through their engagement with dynein. All four function at the centrosome: NUMA1 focuses microtubule minus ends (Merdes et al., 1996; Heald et al., 1996; Merdes et al., 2000; Sikirzhytski et al., 2014; Elting et al., 2014; Hueschen et al., 2017; Aslan et al., 2026), BICD2 tethers the centrosome and the nucleus in G2 and M phases (Splinter et al., 2010; Gallisà-Suñé et al., 2023), NIN nucleates and anchors microtubules (Casenghi et al., 2005; Delgehyr et al., 2005; Redwine et al., 2017), and CCDC88C (also known as Daple) coordinates microtubule dynamics and ciliary positioning (Takagishi et al., 2017; Siletti et al., 2017).

Because this transport occurs during translation, dynein is expected to engage a coiled-coil dimer formed by the nascent polypeptides emerging from separate ribosomes. Co-translational targeting may ensure that an adaptor meets its cargo only at the right place and time, since its C-terminal cargo-binding domain is synthesized last. Alternatively, adaptor dimerization may occur preferentially during translation, which would ensure proper assembly. How translation and dynein-mediated transport are coordinated remains unresolved.

### A new mode of dynein-cargo engagement

Biochemical and structural studies have converged on a unifying framework: dynein cargo adaptors differ in sequence but engage dynein and dynactin through a small number of shared anchor points, with substantial local variation in architecture (Lau et al., 2021; Singh et al., 2024; Teixeira et al., 2025; Aslan et al., 2026; d’Amico et al., 2026). PCNT departs from this framework in two respects. First, its dynein-binding region lies more than 1,000 amino acids from the N-terminus, whereas the dynein-interacting domains of all known cargo adaptors are located at the N-terminus. Second, PCNT is itself the cargo, and its adaptor-like sequences engage and activate dynein without a separate adaptor. This challenges the prevailing view that mammalian dynein relies on a distinct adaptor to engage cargo, and it suggests that dynein recognizes its cargoes in more ways than previously appreciated.

This unusual domain order raises further possibilities. If the N-terminal portion of PCNT binds other proteins, PCNT could also serve as an adaptor that carries those proteins toward the centrosome. In canonical adaptors, the dynein-binding domain is N-terminal and the cargo-binding domain is C-terminal, so the cargo trails the motor. In PCNT, cargo bound N-terminal to the dynein-binding region would lie ahead of dynein, and dynein might therefore push the cargo rather than pull it. Whether PCNT binds and transports other cargoes in this manner, and whether dynein can generate force effectively in this geometry, remain to be tested. More generally, which other cargoes engage dynein and dynactin directly, and how this regulation differs from that of adaptor-mediated transport, remain to be determined.

## Methods

### Constructs

Chimeric PCNT constructs carrying *piggyBac* transposon elements and a doxycycline (Dox)-inducible promoter were generated by replacing all or part of the PCNT sequence in PB-TA-sfGFP-PCNT (854–1960) (Jiang et al., 2021) with synthetic DNA (Twist Bioscience, South San Francisco, CA, USA). In each case, the PB-TA-sfGFP backbone was amplified by PCR with Phusion High-Fidelity DNA Polymerase (F530L, Thermo Fisher Scientific, Waltham, MA, USA) and assembled with the synthetic fragments by isothermal Gibson assembly (Gibson et al., 2009). The constitutively active adaptor construct, sfGFP-tagged human BICD2 (25–423) (BICD2N) (McKenney et al., 2014; Okada et al., 2023), was generated in the same way by replacing the PCNT sequence in PB-TA-sfGFP-PCNT (854–1960) with synthetic DNA encoding BICD2 (25–423). Numbers in parentheses denote amino acid positions.

Constructs for the rapamycin-induced relocalization of peroxisomes were generated as follows. The peroxisome-targeting sequence of human PEX3 (1–42) (Kapitein et al., 2010) was fused to mStay-Gold (mSG1) (Ando et al., 2023) and FK506-binding protein 12 (FKBP). Separately, mScarlet3 (mSL3) (Gadella et al., 2023) was fused to human BICD2 (25–423) and the FKBP12-rapamycin-binding (FRB) domain. Both cassettes were synthesized and cloned between the EcoRI and NotI sites of the lentiviral plasmid pLVX-EF1α-mCherry-N1 (631986, Takara Bio, Mountain View, CA, USA), from which the mCherry sequence had been removed, by Gibson assembly. This yielded pLVX-EF1α-PEX3 (1–42)-mSG1-FKBP and pLVX-EF1α-mSL3-BICD2 (25–423)-FRB, respectively. Additional mSL3- and FRB-tagged adaptor constructs were generated by replacing BICD2 (25–423) in pLVX-EF1α-mSL3-BICD2 (25–423)-FRB, either by Gibson assembly or by site-directed mutagenesis with the Q5 Site-Directed Mutagenesis Kit (E0554S, New England BioLabs, Ipswich, MA, USA).

Constructs for the in vitro dynein motility assays were obtained or generated as follows. The wild-type LIS1 construct (pFastBac backbone, for insect cell expression) and constitutively active human BICD2 (1–598) in the pET28a backbone (for bacterial expression) were gifts from the Reck-Peterson Laboratory (Weill Cornell Medicine), as was the stable HEK293T cell line expressing p62-Halo that was used to purify dynactin. The dynein construct for insect cell expression (Addgene plasmid #111903) was provided by the Carter Laboratory (MRC). PCNT (1393–1780) and PCNT (1393–1910) were amplified by PCR with Q5 High-Fidelity DNA Polymerase (M0491L, New England BioLabs) and substituted for BICD2 (1–598) in the pET28a backbone by Gibson assembly for bacterial expression.

### Cell culture

hTERT-immortalized retinal pigment epithelial (hTERT RPE-1) cells were maintained in Dulbecco’s modified Eagle medium/Ham’s F-12 50/50 mix (DMEM/F-12; 10-092-CV, Corning), and HeLa cells (CCL-2, ATCC, Manassas, VA, USA) were maintained in DMEM (10-017-CV, Corning). Both media were supplemented with 10% fetal bovine serum (FBS; F0926, MilliporeSigma, Burlington, MA, USA) and 1x penicillin-streptomycin-L-glutamine (30-009-CI, Corning). The cells were grown at 37°C in a humidified incubator with 5% CO_2_. The cell lines were not authenticated after they were obtained from their sources, and all tested negative for mycoplasma by PCR with the Universal Mycoplasma Detection Kit (30-1012K, ATCC). None of the cell lines appears on the list of commonly misidentified cell lines maintained by the International Cell Line Authentication Committee.

### Affinity purification of PCNT-dynein complexes

Stable RPE-1 cell lines expressing sfGFP-tagged bait proteins under a Dox-inducible promoter were generated with a *piggyBac* transposon system (Kim et al., 2016; Jiang et al., 2021). After Dox induction, sfGFP or the sfGFP-tagged PCNT and BICD2 baits, together with their associated proteins, were purified by nanobody (Nb)-mediated affinity purification as described previously (Stevens et al., 2024). Approximately 2 × 10^7^ cells from two 15-cm dishes at about 90% confluence were trypsinized, washed twice with phosphate-buffered saline (PBS), and lysed in Nb-lysis buffer [20 mM HEPES-KOH, pH 7.5, 150 mM CH_3_COOK, 2 mM MgSO_4_, 1 mM EGTA, 10% glycerol, 0.1 mM ATP, 0.01% glyco-diosgenin (GDN), 1 mM DTT, 1 mM PMSF, and 1x cOmplete protease-inhibitor cocktail (11873580001, MilliporeSigma)] by Dounce homogenization with a tight pestle (06-435A, Fisher Scientific, Waltham, MA, USA), using approximately 800 µl of buffer per 0.1 g of cell pellet. The homogenates were cleared by centrifugation at 35 000 *g* for 35 minutes at 4°C and incubated for 1 hour at 4°C with magnetic streptavidin beads (75 µl of bead slurry) coated with 37.5 µg of biotinylated anti-GFP Nb. The beads were washed five times with Nb-wash buffer (Nb-lysis buffer containing 0.4% Triton X-100; 1.25 ml per wash) and once with Nb-lysis buffer (1.25 ml). Bound proteins were eluted by incubating the beads for 20 minutes on ice in 75 µl of Nb-elution buffer (50 mM HEPES-KOH, pH 7.5, 200 mM NaCl, 2 mM Mg(CH_3_COO)_2_, 0.5 mM ATP, 0.01% GDN, 0.05% Triton X-100, 1 mM DTT, 1 mM PMSF, and 280 nM SENP^EuB^ protease). The eluates were analyzed by silver staining, Western blotting, and/or mass spectrometry.

### Immunoblotting

Samples were resolved on 4–12% Bolt™ Bis-Tris Plus polyacrylamide gels (NW04125BOX, Thermo Fisher Scientific) and transferred to polyvinylidene difluoride (PVDF) membranes. The membranes were cut horizontally at the 125-kDa marker and blocked for 1 hour at room temperature in Intercept™ Blocking Buffer (927-60001, LI-COR, Lincoln, NE, USA). The upper halves were incubated for 1 hour at room temperature in Intercept™ Blocking Buffer with anti-DYNC1H1 (1:2000, 12345-1-AP, Proteintech, Rosemont, IL, USA) and anti-p150^Glued^ (1:500, 610474, BD Biosciences, San Jose, CA, USA) antibodies, and the lower halves were incubated under the same conditions with an anti-dynein 74-kDa intermediate chain antibody (1:2000, MAB1618, MilliporeSigma) or anti-DCTN6 antibody (1:1000, 16947-1-AP, Proteintech). The membranes were washed once for 15 minutes and three times for 5 minutes each with Tris-buffered saline (20 mM Tris-HCl, pH 7.6, 150 mM NaCl) containing 0.1% Tween-20 (TBS-T), and then incubated for 1 hour at room temperature with IRDye® secondary antibodies (1:30,000, LI-COR) in Intercept™ Blocking Buffer. After the same series of TBS-T washes, the membranes were dried and imaged on an Odyssey CLx Infrared Imaging System (Model 9140, LI-COR).

### Mass spectrometry

#### Sample preparation

Samples were exchanged into digestion buffer by a modified SP3 protocol (Hughes et al., 2019). Approximately 250 µg of Cytiva SpeedBead magnetic carboxylate-modified particles (65152105050250 and 4515210505250, Cytiva, Marlborough, MA, USA), mixed at a 1:1 ratio, was added to each sample, and 100% ethanol was added to a final concentration of at least 50%. The samples were incubated with gentle shaking for 15 minutes and washed three times with 80% ethanol. Proteins were eluted from the beads in 200 mM EPPS, pH 8.5, containing Lys-C (129-02541, Wako, Osaka, Japan) and digested overnight at room temperature with vigorous shaking. Trypsin (90305, Thermo Fisher Scientific) was then added, and the digestion was continued for 6 hours at 37°C.

Acetonitrile was added to each sample to a final concentration of approximately 33%, and the peptides were labeled in the presence of the SP3 beads with approximately 62.5 µg of TMTpro reagents (18 channels in total; A44520 and A52045, Thermo Fisher Scientific). After labeling efficiency was confirmed to exceed 97%, excess TMT was quenched with hydroxylamine at a final concentration of 0.3%. The samples were pooled, and acetonitrile was removed by vacuum centrifugation. The pooled sample was acidified with formic acid, desalted on a StageTip, eluted into autosampler inserts (6PSV9-03FIVPT, Thermo Fisher Scientific), dried in a vacuum concentrator, and reconstituted in 5% acetonitrile and 5% formic acid for LC-MS/MS.

#### Liquid chromatography and tandem mass spectrometry

Data were collected on an Orbitrap Eclipse mass spectrometer coupled to a Proxeon NanoLC-1200 UH-PLC (Thermo Fisher Scientific). The 100-µm capillary column was packed in-house with 35 cm of Accucore 150 resin (2.6 µm, 150 Å; Thermo Fisher Scientific), and each run lasted 180 minutes. A FAIMS device was used with compensation voltages of −40 V, −60 V, and −80 V (Schweppe et al., 2019). MS1 scans were acquired in the Orbitrap (resolution, 60,000; scan range, 400–1600 Th; automatic gain control (AGC) target, 4 × 10^10^; maximum ion injection time, automatic). MS2 scans were acquired in the Orbitrap after higher-energy collision dissociation (HCD; resolution, 50,000; AGC target, 250%; normalized collision energy, 36; isolation window, 0.5 Th; maximum ion injection time, 86 ms).

#### Data analysis

Spectra were searched against all entries of the human UniProt database (downloaded in May 2021) concatenated with a decoy database of the same sequences in reverse order (Elias and Gygi, 2007). Raw files were converted to mzXML, and monoisotopic peaks were reassigned with Monocle (Rad et al., 2021). Searches were performed with Comet (Eng et al., 2013) using a precursor ion tolerance of 50 ppm and a fragment bin tolerance of 0.02. TMTpro labels on lysine residues and peptide N-termini (+304.207 Da) and carbamidomethylation of cysteine residues (+57.021 Da) were set as static modifications, and oxidation of methionine residues (+15.995 Da) was set as a variable modification. Peptide-spectrum matches (PSMs) were filtered to a 1% false discovery rate (FDR) with a linear discriminant, and proteins were assembled to a final protein-level FDR of 1% (Huttlin et al., 2010).

TMT reporter ion intensities were measured in a 0.003-Da window around the theoretical *m/z* of each reporter ion and corrected for the isotopic impurities of the TMT reagents according to the manufacturer’s specifications. Peptides were excluded if their summed signal-to-noise ratio (S/N) across all TMT channels was below 180 or if their precursor isolation specificity was below 0.5. Protein abundance was calculated by summing the S/N values of all peptides assigned to a protein, and only proteins identified by at least two peptides were retained. Abundances were normalized to that of the PCNT segment shared by GFP-PCNT (854–1960) and GFP-PCNT (854–1960)Δ1393–1525 baits (residues 854–1393 and 1525–1960). Normalized abundances were compared by Welch’s two-sample *t*-test, which does not assume equal variances, and the raw *P* values were adjusted by the Benjamini-Hochberg (BH) procedure (Benjamini and Hochberg, 1995) to control the FDR. Proteins with a BH-adjusted *p*-value (*q*-value) below 0.03 and a fold change of at least 2 were considered significantly depleted or enriched; no more than 3% of these were expected to be false positives.

### Protein expression and purification

#### Dynein and LIS1 expression

Human dynein and LIS1 were expressed in Sf9 cells grown in Sf-900 serum-free medium (10902096, Thermo Fisher Scientific) as described (Siva et al., 2026). The pACEBac1 plasmid encoding human dynein and the pFastBac plasmid encoding wild-type LIS1 were transformed into competent DH10EMBacY cells by heat shock at 42°C for 15 seconds, followed by recovery for 6 hours at 37°C with shaking at 220 rpm in S.O.C. medium (15544034, Thermo Fisher Scientific). The cells were plated on LB-agar containing tetracycline (10 µg/ml), gentamicin (7 µg/ml), kanamycin (50 µg/ml), IPTG (40 µg/ml), and Bluo-Gal (100 µg/ml), and colonies containing the plasmid of interest were identified by blue/white screening after 48–72 hours of growth. For dynein, white colonies were tested by PCR for the presence of all dynein genes. Selected colonies were grown for 14–16 hours in LB medium containing tetracycline (10 µg/ml), gentamicin (7 µg/ml), and kanamycin (50 µg/ml) at 37°C with shaking at 220 rpm, and bacmid DNA was isolated by isopropanol extraction.

For each bacmid, approximately 1 × 10^6^ Sf9 cells in a well of a six-well plate were transfected with 1–2 µg of purified bacmid using FuGENE HD transfection reagent (E2311, Promega, Madison, WI, USA) according to the manufacturer’s instructions. The cells were incubated without agitation at 27°C in a humidified incubator for 3 days, and the viral supernatant (V0) was collected by centrifugation at 1000 *g* for 5 minutes at 4°C. To amplify the virus, 1 ml of V0 was used to infect 50 ml of Sf9 cells at 1 × 10^6^ cells/ml.

The infected culture was incubated at 27°C with shaking at 105 rpm for 3 days, and the supernatant (V1) was harvested by centrifugation as above. For protein expression, 4 ml of V1 was used to infect 400 ml of Sf9 cells at 1 × 10^6^ cells/ml, and the culture was incubated under the same conditions for 3 days. The cells were pelleted by centrifugation at 2100 *g*, washed with 10 ml of cold PBS, flash-frozen in liquid nitrogen, and stored at −80°C until purification.

##### Dynein purification

Human dynein was purified as described (Siva et al., 2026). All steps were carried out at 4°C unless stated otherwise. Each cell pellet was resuspended in 40 ml of dynein-lysis buffer (50 mM HEPES-KOH, pH 7.4, 100 mM NaCl, 1 mM DTT, 0.1 mM Mg-ATP, and 10% (v/v) glycerol) supplemented with 0.2 mM AEBSF (A-540-1, GoldBio, St. Louis, MO, USA), 81 µM bestatin (AAJ60384-MB, VWR, Radnor, PA, USA), 7 µM E-64 (E-064-25, GoldBio), 21 µM leupeptin (L-010-100, GoldBio), 7 µM pepstatin A (78781-879, VWR), 1 mM benzamidine (12072-10G, MilliporeSigma), and 1 mM PMSF (10837091001, MilliporeSigma). The cells were lysed with a Dounce homogenizer, and the lysate was clarified by centrifugation at 183 960 *g* for 88 minutes in a Type 70 Ti rotor (337922, Beckman Coulter, Brea, CA, USA).

The clarified supernatant was incubated for 4 hours with rotation with 2 ml of IgG Sepharose 6 Fast Flow beads (17015901, Cytiva) equilibrated in dynein-lysis buffer. The beads were collected in a gravity column and washed with 200 ml of dynein-lysis buffer and 300 ml of tobacco etch virus (TEV) buffer (50 mM Tris-HCl, pH 8.0, 250 mM CH_3_COOK, 2 mM Mg(CH_3_COO)_2_, 1 mM EGTA, 1 mM DTT, 0.1 mM Mg-ATP, and 10% (v/v) glycerol). For fluorescent labeling of the SNAP tag, the bead-bound dynein was incubated with 5 µM SNAP-JF646 (Lavis Laboratory, Janelia) for 10 minutes at room temperature. The beads were washed again with 300 ml of TEV buffer, resuspended in 15 ml of TEV buffer containing 0.05 mg/ml TEV protease, and incubated overnight with gentle rotation. The cleaved protein was collected in the flow-through of a gravity column and concentrated to 500 µl in a 100-kDa MWCO concentrator (UFC210024, MilliporeSigma).

The concentrate was injected onto a TSKgel G4000SWxl size-exclusion column (0008542, Tosoh Bioscience, King of Prussia, PA, USA) equilibrated in GF150 buffer (25 mM HEPES-KOH, pH 7.4, 150 mM KCl, 1 mM MgCl_2_, and 1 mM DTT) supplemented with 0.1 mM Mg-ATP, and the column was run at 0.75 ml/min. The peak fractions were collected, exchanged into GF150 buffer containing 0.1 mg/ml ATP and 10% glycerol, concentrated to 0.1–0.5 mg/ml, flash-frozen in small aliquots, and stored at −80°C.

##### LIS1 purification

LIS1 was purified from Sf9 cells as described for dynein, except that LIS1-lysis buffer (30 mM HEPES-KOH, pH 7.4, 50 mM CH_3_COOK, 2 mM Mg(CH_3_COO)_2_, 1 mM EGTA, 300 mM KCl, 1 mM DTT, and 10% (v/v) glycerol) was used in place of dynein-lysis buffer. The LIS1-lysis buffer was supplemented with 0.2 mM AEBSF, 81 µM bestatin, 7 µM E-64, 21 µM leupeptin, 7 µM pepstatin A, 1 mM benzamidine, and 1 mM PMSF. After lysis, centrifugation, and binding to IgG Sepharose beads, the beads were collected in a gravity column and washed with 20 ml of LIS1-lysis buffer, 200 ml of LIS1-TEV buffer (10 mM Tris-HCl, pH 8.0, 150 mM CH_3_COOK, 2 mM Mg(CH_3_COO)_2_, 1 mM EGTA, 1 mM DTT, and 10% (v/v) glycerol) supplemented with 100 mM CH_3_COOK, and 100 ml of LIS1-TEV buffer. The beads were then incubated overnight with 0.05 mg/ml TEV protease. The cleaved protein was collected in the flow-through of a gravity column and concentrated to 500 µl in a 30-kDa MWCO concentrator (UFC803008, Millipore-Sigma).

The concentrate was injected onto a Superose 6 Increase 10/300 GL column (29091596, Cytiva) and eluted at 0.75 ml/min with GF150 buffer supplemented with 10% (v/v) glycerol. The peak fractions were collected, concentrated to 0.2–1 mg/ml in a 30-kDa MWCO concentrator, flash-frozen in small aliquots, and stored at −80°C.

##### Dynactin purification

Dynactin was purified as described (Redwine et al., 2017). HEK293T cells stably expressing p62-Halo-3xFLAG were grown in suspension to 1.5 × 10^6^ cells/ml and harvested by centrifugation at 2100 *g* for 15 minutes. Frozen pellets from 3 L of culture, stored at −80°C, were resuspended in 80 ml of dynactin-lysis buffer (30 mM HEPES-KOH, pH 7.4, 50 mM CH_3_COOK, 2 mM Mg(CH_3_COO)_2_, 1 mM EGTA, 1 mM DTT, and 10% (v/v) glycerol) supplemented with 0.5 mM Mg-ATP, 0.2% Triton X-100, 0.2 mM AEBSF, 81 µM bestatin, 7 µM E-64, 21 µM leupeptin, 7 µM pepstatin A, 1 mM benzamidine, and 1 mM PMSF. The lysate was clarified by centrifugation at 66 000 *g* for 30 minutes at 4°C in a Type 70 Ti rotor. The supernatant was incubated overnight at 4°C with rotation with 1.5 ml of anti-FLAG M2 affinity gel (A2220, MilliporeSigma). The beads were collected in a gravity column and washed with at least 50 ml of wash buffer (dynactin-lysis buffer supplemented with mM Mg-ATP and 0.02% Triton X-100), 100 ml of wash buffer supplemented with 250 mM CH_3_COOK, and 100 ml of wash buffer. Dynactin was eluted by incubating the beads with 1 ml of elution buffer (wash buffer containing 2 mg/ml 3xFLAG peptide).

The eluate was diluted to 2 ml in Buffer A (50 mM Tris-HCl, pH 8.0, 2 mM Mg(CH_3_COO)_2_, 1 mM EGTA, and 1 mM DTT) and loaded at 1 ml/min onto a MonoQ 5/50 GL column (17-5166-01, Cytiva). Bound protein was eluted with a linear gradient from 35% to 100% Buffer B (50 mM Tris-HCl, pH 8.0, 1 M CH_3_COOK, 2 mM Mg(CH_3_COO)_2_, 1 mM EGTA, and 1 mM DTT) over 26 column volumes. The fractions that eluted between 75% and 80% Buffer B were pooled, exchanged into GF150 buffer containing 10% glycerol, concentrated to 0.02–0.1 mg/ml, flash-frozen in small aliquots, and stored at −80°C.

#### Adaptor and PCNT expression

Adaptor and PCNT constructs carrying N-terminal HaloTags were expressed in bacteria and purified as described (Siva et al., 2026). Each construct was transformed into competent BL21(DE3) cells (C2527H, New England Biolabs), and the cells were grown in 1.5 L of LB medium at 37°C with shaking until the OD_600_ reached 0.4–0.6. Protein expression was induced with 0.1 mM IPTG, and the cultures were grown overnight at 16°C. The cells were pelleted by centrifugation, flash-frozen, and stored at −80°C until use.

#### Adaptor and PCNT purification

All steps were carried out at 4°C unless stated otherwise. Frozen pellets from 1.5 L of culture were resuspended in 40 ml of adaptor-lysis buffer (30 mM HEPES-KOH, pH 7.4, 50 mM CH_3_COOK, 2 mM Mg(CH_3_COO)_2_, 1 mM EGTA, 1 mM DTT, and 10% (v/v) glycerol) supplemented with mM AEBSF, 81 µM bestatin, 7 µM E-64, 21 µM leupeptin, 7 µM pepstatin A, 1 mM benzamidine, 1 mM PMSF, and 1 mg/ml lysozyme. The suspension was incubated on ice for 30 minutes and sonicated, and the lysate was clarified by centrifugation at 66 000 *g* for 30 minutes in a Type 70 Ti rotor.

The supernatant was incubated for 2 hours with rotation with 2 ml of IgG Sepharose 6 Fast Flow beads equilibrated in adaptor-lysis buffer. The beads were collected in a gravity column and washed with adaptor-lysis buffer supplemented with 150 mM CH_3_COOK and with 50 ml of adaptor-TEV buffer (50 mM Tris-HCl, pH 8.0, 150 mM CH_3_COOK, 2 mM Mg(CH_3_COO)_2_, 1 mM EGTA, 1 mM DTT, and 10% (v/v) glycerol). The washed beads were then incubated overnight with rotation in 15 ml of adaptor-TEV buffer containing up to 0.2 mg/ml TEV protease. The cleaved protein was collected, concentrated to 1 ml in a 30-kDa MWCO concentrator, filtered through an Ultrafree-MC VV filter (UFC30GV0S, MilliporeSigma), and diluted to 2 ml in Buffer A (30 mM HEPES-KOH, pH 7.4, 50 mM CH_3_COOK, 2 mM Mg(CH_3_COO)_2_, 1 mM EGTA, 10% (v/v) glycerol, and 1 mM DTT).

The diluted protein was injected at 1 ml/min onto a 1-ml HiTrap Q column (17115301, Cytiva) or a MonoQ 5/50 GL column (17-5166-01, Cytiva) and eluted with a linear gradient from 0 to 100% Buffer B (30 mM HEPES-KOH, pH 7.4, 1 M CH_3_COOK, 2 mM Mg(CH_3_COO)_2_, 1 mM EGTA, 10% (v/v) glycerol, and 1 mM DTT) over 26 column volumes. The peak fractions containing HaloTag-tagged protein were pooled, concentrated, diluted to 0.5 ml in GF150 buffer, and injected onto a Superdex 200 10/300 column (17517501, Cytiva) equilibrated in GF150 buffer and run at 0.5 ml/min. The peak fractions were collected and exchanged into GF150 buffer containing 10% glycerol.

#### Adaptor and PCNT labeling

HaloTag-tagged adaptors and PCNT were labeled by incubating the concentrated protein fractions with a 10-fold molar excess of HaloTag ligand conjugated to TMR or JF549 (GA1110, Promega) for 10 minutes at room temperature. Excess dye was removed on a Micro Bio-Spin desalting column pre-equilibrated with GF150 buffer containing 10% (v/v) glycerol. The labeled proteins were flash-frozen in small aliquots and stored at −80°C.

### Evolutionary signature analysis

Evolutionary signatures of protein segments were calculated with the established pipeline (Pritišanac et al., 2026) (code at https://github.com/IPritisanac/IDR_ES). One-to-one orthologs of the selected proteins were collected from the OMA database through OmaDB (Kaleb et al., 2019) and aligned with MAFFT (Katoh and Standley, 2013), and the sequences aligning with each segment of interest were extracted as described (Pritišanac et al., 2026). The evolutionary signature of a segment was defined as the set of z-scores comparing the distribution of each of 150 sequence features in its orthologous set with the distribution of features in 1000 simulated ortholog sets, each of which was generated by simulated evolution as described (Pritišanac et al., 2026). The magnitude of the z-scores was capped at 10. Two features, “aromatic_spacing_meanZ” and “omega_aromatic_meanZ”, were excluded because some segments contain no aromatic residues.

The signatures were clustered by hierarchical agglomerative clustering with average linkage and cosine distance, applied to both rows and columns, using the plotly clustergram (https://dash.plotly.com/dash-bio/clustergram). Twelve segments were excluded because they were too short or their ortholog alignments were poor: Q32MZ4 (128–250), Q9HCU9 (51–98), N-SPDL1 (1–22), N-RAB45 (1–20), O60239 (97–145), N-JIP3 (1–11), N-NIN (1–7), B-JIP3 (12–386), C-CDR2L (444–465), C-RAB11FIP3 (713–756), C-JIP3 (387–1336), and Q9UHQ4 (166–233).

### Synthetic protein design

Synthetic proteins (SPs) were designed to match the bulk sequence features of PCNT (residues 1393–1525; referred to as PCNT in this section) using a published algorithm (Strome et al., 2023). The algorithm starts from a user-defined or random sequence, introduces point mutations iteratively, and greedily retains those that bring the feature vector of the design closer to that of PCNT. The distance between the two vectors is the Euclidean distance:

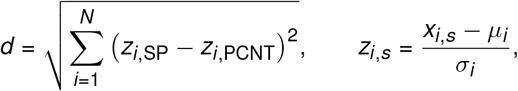

where *N* is the number of features, *x*_*i, s*_ is the raw value of feature *i* for sequence *s*, and *µ*_*i*_ and *σ*_*i*_ are the mean and standard deviation of feature *i* across the human IDRome.

We initialized SP1^ING4^ and SP5^STAT1^ with the UniProt-annotated predicted coiled-coil regions of ING4 (Q9UNL4, residues 25–118) and STAT1 (P42224, residues 136–317), respectively, and SP2, SP3, and SP4 with random sequences in which each amino acid was drawn with equal probability. A Colab notebook for the design algorithm is available at https://colab.research.google.com/drive/ 15cX0QzDVm-owJgIk0AgZIARyXf1NpNCx and a web-based version of the method is also available (Huang et al., 2026). For each SP, we selected one design at random from 3–15 candidates for functional testing.

### Coiled-coil predictions

Coiled-coil content was predicted for the isolated sequences of ING4 (25–118), SP1^ING4^, SP2–SP4, SP5^STAT1^, and PCNT (1393–1525) with Ncoils (Lupas et al., 1991) and four coiled-coil predictors run on the MPI Bioinformatics Toolkit (Gabler et al., 2020). Ncoils was run with default weights, matrix, and settings, including a window size of 21 and a probability threshold of *≥* 0.5; residues above the threshold were counted as coiled-coil. PCOILS (Gruber et al., 2006) was run with the MTIDK matrix and a 21-residue window, and residues with a probability *≥* 0.5 were counted. MARCOIL (Delorenzi and Speed, 2002) was run with its default calibration, and residues with a per-residue coiled-coil probability *≥* 50% were counted. DeepCoil (Ludwiczak et al., 2019) assigns a probability to each residue, and residues with a probability *≥*0.5 were counted. For DeepCoil2 (Ludwiczak et al., 2019), residues within the segments that the program called as coiled-coil were counted. For each construct and predictor, coiled-coil content was the number of counted residues divided by the sequence length. Because such predictors tend to overestimate or underestimate the extent of coiled-coil regions (Simm et al., 2021), we considered the output of these five predictors and reported the mean and the sample standard deviation (SD, *n −* 1 denominator) of the five values (**Extended Data Table 2**). This SD reflects disagreement between predictors, not experimental variability.

Coiled-coil content of Δ1393–1525 was set to zero, because the predicted region is deleted.

### Structural predictions

Structures were predicted with the ColabFold implementation (Mirdita et al., 2022) of AlphaFold2 (Evans et al., 2021; Jumper et al., 2021). Models were generated with ColabFold v1.6.1 and AlphaFold-Multimer v3 weights, using 12 recycles per model and four random seeds, and no structural templates were used. The PCNT-DHC complex in **Fig. 4b** was modeled with two copies of PCNT residues 1397–1517 (UniProt O95613) and two copies of DHC residues 540–910 (UniProt Q14204), corresponding to one PCNT dimer and one DHC dimer. The PCNT-2DHC complex in **Fig. 4c** was modeled with two copies of PCNT residues 1397–1517 and four copies of DHC residues 540–910 (two DHC dimers) using the same parameters.

Predicted aligned error (PAE) values relative to DHC residues Y^827^ and R^759^ were mapped onto the predicted structures with a custom Python script adapted from PointPAE (Chaaban and Carter, 2022) (https://github.com/sami-chaaban/PointPAE; https://doi.org/10.5281/zenodo.6792801).

For each reference residue, the script extracted the PAE values along both axes of the PAE matrix, averaged them, and wrote a ChimeraX attribute file for visualization. Structures were visualized in UCSF ChimeraX (Goddard et al., 2018; Pettersen et al., 2021; Meng et al., 2023).

### Condensate motility assay and analysis

#### Data collection

The assay was performed as described previously (Jiang et al., 2021, 2022). RPE-1 cells stably expressing GFP-tagged PCNT constructs under a Dox-inducible promoter were seeded in 35-mm glass-bottom dishes (D35C4-20-1.5-N, Cellvis, Mountain View, CA, USA) and incubated with 1 µg/ml Dox to induce protein expression and condensate formation. The cells were imaged live in a humidified chamber with 5% CO_2_ at 37°C (wrap-around environmental incubator, Okolab, Pozzuoli, Italy). Images were acquired on a spinning disk confocal microscope (Dragonfly 620SR, Andor Technology, Belfast, UK) with a 63x/1.40 NA (numerical aperture) HC PL APO objective (Leica, Wetzlar, Germany). Acquisition was controlled by Fusion software (Andor Technology), and images were captured with an iXon Ultra 888 electron-multiplying CCD (EMCCD) camera.

#### Analysis

Image segmentation, object tracking, and quantification were performed with the Python-based TRACES workflow (Jiang et al., 2022). Confocal z-stacks were projected by maximum intensity and split into channels for GFP-PCNT (854–1960) (condensates), mScarlet-i-H2A (nuclei), and miRFP670nano3-CETN2 (centrosomes). Intensity and size thresholds were applied to each channel to identify the objects of interest. The Analyze Particles function in Fiji (Schindelin et al., 2012) was used to assign each object an identification number and to measure its center-of-mass coordinates (*x, y*) and projected area. The Fiji measurements were then processed by TRACES to track the objects in each cell and to calculate the distance of each condensate to the centrosome over a 200-minute imaging period. Because phase separation began asynchronously, the time courses were aligned to the onset of phase separation of each condensate (time 0). Data were processed with Pandas (Reback et al., 2020) and NumPy (Harris et al., 2020), and plotted with Matplotlib (Hunter, 2007) and GraphPad Prism 11 (GraphPad, San Diego, CA, USA).

#### Statistic

Statistical analyses were performed in Prism 11 (GraphPad). The distance between each condensate and the centrosome was log_2_-transformed, and a linear regression of log_2_ distance against time was fitted for each construct. Slopes were compared pairwise by *F*-test, with the null hypothesis that the two regression lines have equal slopes. Each construct was compared with the PCNT (854–1960) and PCNT (854–1960)Δ1393–1525 controls, as indicated in **Fig. 1b, Fig. 3**, and **Extended Data Table 3**.

### Peroxisome motility assay and analysis

*Data collection*. HeLa cells were seeded at 70% confluence in 35-mm glass-bottom dishes and transfected with plasmids encoding PEX3-mSG1-FKBP and either mSL3-BICD2N-FRB, mSL3-PCNT (1393–1910)-FRB, or a BICD2N or PCNT variant, using TransIT®-LT1 transfection reagent (MIR 2304, Mirus Bio, Madison, WI, USA) according to the manufacturer’s protocol. After 24 hours, the cells were treated with DMSO or 0.5 µM rapamycin and imaged immediately for up to 20 minutes on a spinning disk confocal microscope (Dragonfly 620SR, Andor Technology) with a 63x/1.40 NA or 100x/1.40 NA objective.

#### Analysis

Confocal z-stacks were projected by maximum intensity and the background was subtracted using the rolling-ball method with the sliding paraboloid option in Fiji. A region of interest (ROI) was drawn around the boundary of each cell. Within the ROI, the area covered by peroxisomes (mSG1) and the area of the entire cell (mSL3) were each defined as the area containing 90% of the total fluorescence in that channel, using a custom ImageJ macro. The macro ranked the pixels by intensity in descending order, determined the smallest number of the brightest pixels that accounted for 90% of the integrated fluorescence, and expressed this number as a fraction of the pixels in the ROI. For each cell at each time point, a peroxisome-to-total cell area (P/T) ratio was calculated by dividing the 90% fluorescence area of mSG1 by that of mSL3. Peroxisome redistribution was expressed as the peroxisome redistribution index (PRI), the fractional change in the P/T ratio after rapamycin addition:

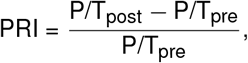

where P/T_pre_ was measured 2 minutes before and P/T_post_ 15 minutes after rapamycin addition. Negative PRI values indicated clustering of peroxisomes relative to the distribution of the adaptor, and positive values indicated dispersion.

#### Statistics

Statistical analyses were performed in Prism 11 (GraphPad) and consisted of three tests. First, a two-tailed paired *t*-test compared the P/T ratios before and after rapamycin addition to determine whether peroxisomes were trafficked. Second, a one-sample *t*-test against a hypothetical mean of zero determined whether each construct changed the distribution of peroxisomes from baseline. Third, mutant constructs were compared with their respective controls by two one-way ANOVAs, each followed by Dunnett’s multiple comparisons test. Within the BICD2 series, the mutants mtHBS1-BICD2N and mtSPDLY-BICD2N were compared with BICD2N. Within the PCNT series, PCNT (1393–1525), PCNT (1393–1780), mtN-HBS1, mtC-HBS1, and mtSPDLY were compared with PCNT (1393–1910).

### Single-molecule TIRF microscopy

Single-molecule imaging was performed on an inverted microscope (Ti2-E Eclipse, Nikon) with a TIRF 100x/1.49 NA APO objective (MRD01991, Nikon, Melville, NY, USA) and a LUNF-XL laser launch (Nikon) providing 405-, 488-, 561-, and 640-nm lasers. The excitation path was filtered through a quad bandpass filter cube (89901 ET, Chroma, Bellows Falls, VT, USA), and emission was filtered through band-specific filters (Chroma) in a high-speed filter wheel (Finger Lakes Instrumentation, Lima, NY, USA) and detected with an iXon Ultra 897 EMCCD camera (Andor Technology). The microscope and image acquisition were controlled by NIS-Elements Advanced Research software (Nikon).

### Single-molecule motility assay and analysis

#### Data collection

Flow chambers were assembled as described previously (Case et al., 1997; Siva et al., 2026) with No. 1½ coverslips (Corning, Corning, NY, USA) functionalized with PEG-biotin. Microtubules were prepared by polymerizing 20 µM bovine tubulin (~10% biotin-tubulin and ~10% Alexa-488 tubulin) at 37°C for 30 minutes and stabilizing them with 20 µM taxol. The chambers were incubated for 3 minutes with 1 mg/ml streptavidin in assay buffer (30 mM HEPES-KOH, pH 7.4, 2 mM Mg(CH_3_COO)_2_, 1 mM EGTA, 10% glycerol, 1 mM DTT, and 20 µM taxol) and washed twice with 20 µl of assay buffer. Taxol-stabilized microtubules (final concentration ~ 0.1–0.4 µM) were then flowed in and incubated for 3 minutes, and the chambers were washed twice with assay buffer containing 1 mg/ml casein.

Dynein-dynactin-adaptor/PCNT complexes were assembled by mixing dynein (0.75 nM final concentration), dynactin, LIS1, and adaptor or PCNT at a molar ratio of 1:2:40:20 and incubating the mixture for 10 min on ice. The complexes were diluted in assay buffer supplemented with 90 mM CH_3_COOK, 0.45% methyl-cellulose, 1 mg/ml casein, and 0.7% Pluronic F127 and incubated for another 10 min on ice. Immediately before imaging, the samples were mixed with assay buffer to a final composition of 36.75 mM KCl, 83.25 mM CH_3_COOK, 0.4% methylcellulose, 0.5% Pluronic F127, 1 mg/ml casein, 71.5 mM β-mercaptoethanol, mg/ml glucose catalase, 1.2 mg/ml glucose oxidase, 0.4% glucose, and 2.5 mM Mg-ATP, and introduced into the flow chamber.

The microtubules were imaged first as a single snapshot of the field of view. Dynein (labeled with JF646) and adaptor or PCNT [labeled with tetramethylrhodamine (TMR) for BICD2 or JF549 for PCNT] were then imaged every 300 ms for 2 minutes. The microtubule channel was imaged again at the end of each movie to assess stage drift, and movies showing drift were excluded.

#### Analysis

Kymographs were generated from the movies with ImageJ macros as described (Roberts et al., 2014) and analyzed with KymoButler (Jakobs et al., 2019). The landing rate was determined for each microtubule as

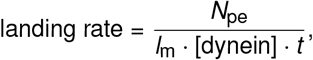

where *N*_pe_ is the number of processive events, *l*_m_ is the length of the microtubule (in µm), and *t* is the length of the movie (in minute). Velocity (µm/s) and run length (µm) were measured from the kymographs for the same processive events. Statistical analyses were performed in Prism 11 (GraphPad).

#### Reproducibility

Data were collected with two independent preparations of dynein and of each PCNT construct. Because the density of microtubules on the coverslip can influence the landing rate, the chambers were inspected visually and used only if their microtubule densities were similar.

## Contributions

W.Z., A.M.M., and L.-E.J. conceived and planned the project. W.Z., A.G.S., T.T.N., J.P.G., S.M., J.-L.S., T.H.H., A.G.I., X.J., J.M., H.A., M.E.D., A.M.M., and L.-E.J. generated reagents, performed experiments, and/or collected and analyzed data. A.G.S. and A.M.M. wrote codes for evolutionary signature analyses and designed synthetic proteins. L.-E.J. wrote the manuscript with input from all authors. All authors reviewed and edited the manuscript.

## Acknowledgments

We thank all members of the Jao Lab for helpful discussions. The authors also thank Dr. Richard McKenney for insightful feedback on the project. Tandem mass tag mass spectrometry was performed by the Thermo Fisher Scientific Center for Multiplexed Proteomics (TCMP) at Harvard Medical School (https://tcmp.hms.harvard.edu). We thank Drs. Julian Mintseris and Jonathan Van Vranken at TCMP for outstanding assistance in performing and analyzing mass spectrometry data. Experiments were performed in part through the use of the UC Davis Campus Shared Flow Cytometry Resource, the SUNY Upstate Medical University Research Flow Core, and Blatt BioImaging Center at Syracuse University. Molecular graphics and analyses performed with UCSF ChimeraX, developed by the Resource for Biocomputing, Visualization, and Informatics at the University of California, San Francisco, with support from National Institutes of Health R01-GM129325 and the Office of Cyber Infrastructure and Computational Biology, National Institute of Allergy and Infectious Diseases. The manuscript was formatted using the LaTeX template developed by Stephen Royle and Ricardo Henriques Labs.

## Extended Data Information

**Extended Data Figure 1.**
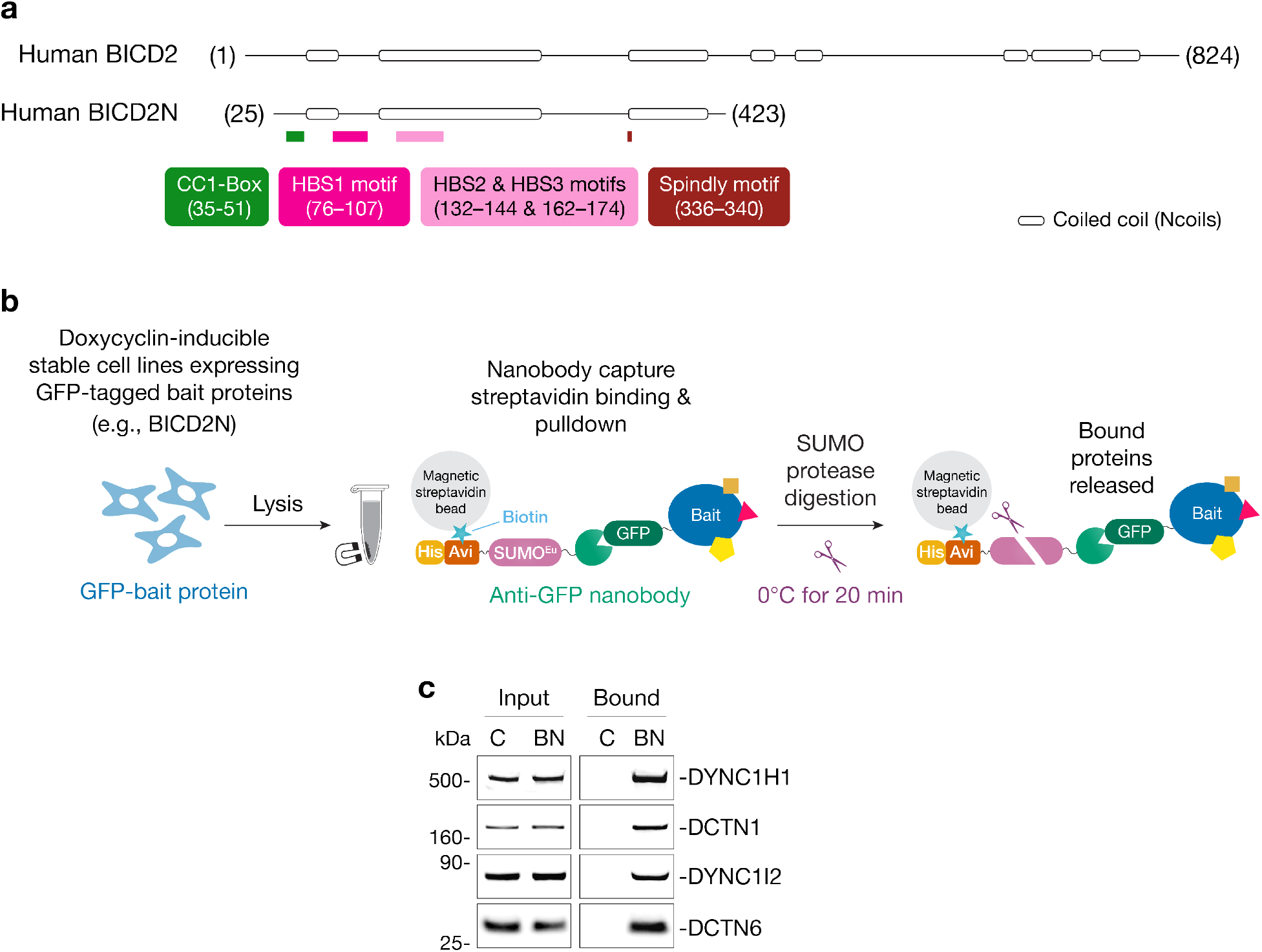
Nanobody-mediated affinity purification of dynein-dynactin-BICD2N complexes from RPE-1 cells. **a**, Domain structures of full-length human BICD2 and its N-terminal fragment (residues 25–423). Locations are indicated for the CC1 box (dynein light intermediate chain, DLIC, binding), the HBS1, HBS2, and HBS3 motifs (dynein heavy chain, DHC, binding), and the Spindly motif (dynactin pointed-end binding). Coiled-coil regions were predicted with Ncoils (Lupas et al., 1991). **b**, Schematic of the anti-GFP nanobody-mediated purification protocol (adapted from Stevens et al., 2024). **c**, Anti-GFP immunoprecipitates from cells expressing GFP-FLAG (C) or GFP-BICD2N-FLAG (BN) were analyzed by immunoblotting for dynein heavy chain (DYNC1H1), dynein intermediate chain (DYNC1I2), and the dynactin subunits DCTN1 and DCTN6. Similar results were obtained in more than three biological replicates.

**Extended Data Figure 2.**
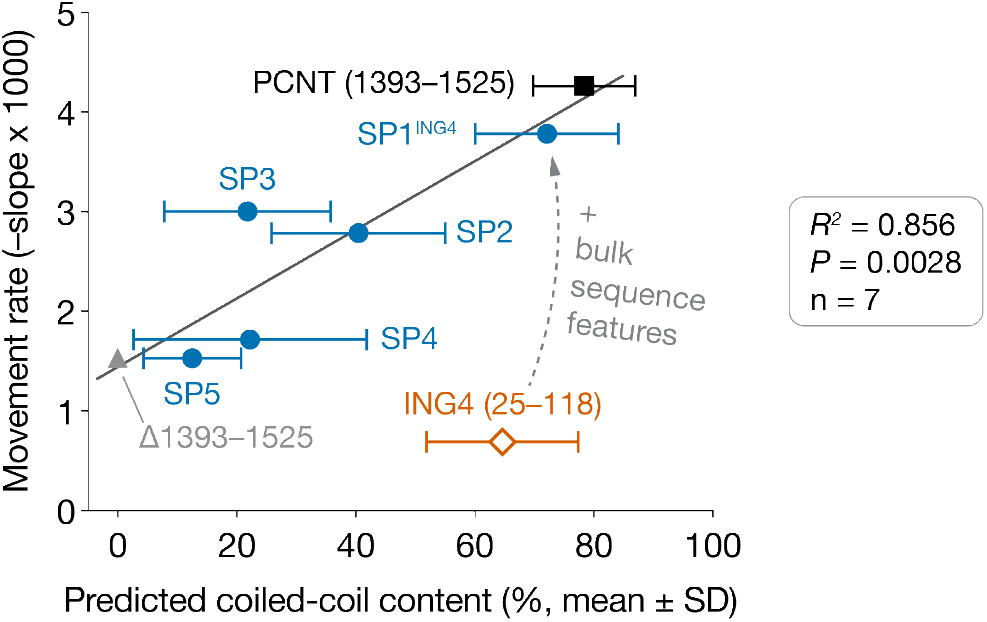
Predicted coiled-coil content of the synthetic sequences correlates with their ability to restore dynein-mediated movement of PCNT condensates. Movement rate, *−* (slope *×*1000), plotted against predicted coiled-coil content (%, mean *±* SD of five predictors) for PCNT (854–1960) constructs in which residues 1393–1525 were retained (PCNT (1393–1525); black square), deleted (Δ1393–1525; gray triangle), or replaced with one of five synthetic proteins (SP1^ING4^, SP2, SP3, SP4, and SP5; blue circles) or with ING4 (25–118) (orange diamond). Movement rate is the negative of the slope of the linear fit of log_2_ condensate-to-centrosome distance against time (**Fig. 3** and **Extended Data Table 3**); higher values indicate faster movement. Coiled-coil content is the percentage of residues predicted to be in a coiled-coil, averaged over five predictors (Ncoils, PCOILS, MARCOIL, DeepCoil, and DeepCoil2; **Extended Data Table 2**). Horizontal bars show the SD across predictors, not experimental variability. Coiled-coil content of Δ1393–1525 was set to zero because the region is deleted. The line is a linear regression fitted to the seven constructs other than ING4 (*R*^2^ = 0.856, *n* = 7; *P* = 0.0028, two-tailed *t*-test of the correlation coefficient with *n −* 2 degrees of freedom). ING4 (open diamond) has high predicted coiled-coil content but little movement and was not included in the fit. The dashed arrow indicates that SP1^ING4^ was derived from ING4 by introducing the bulk sequence features of PCNT (1393–1525).

**Extended Data Figure 3.**
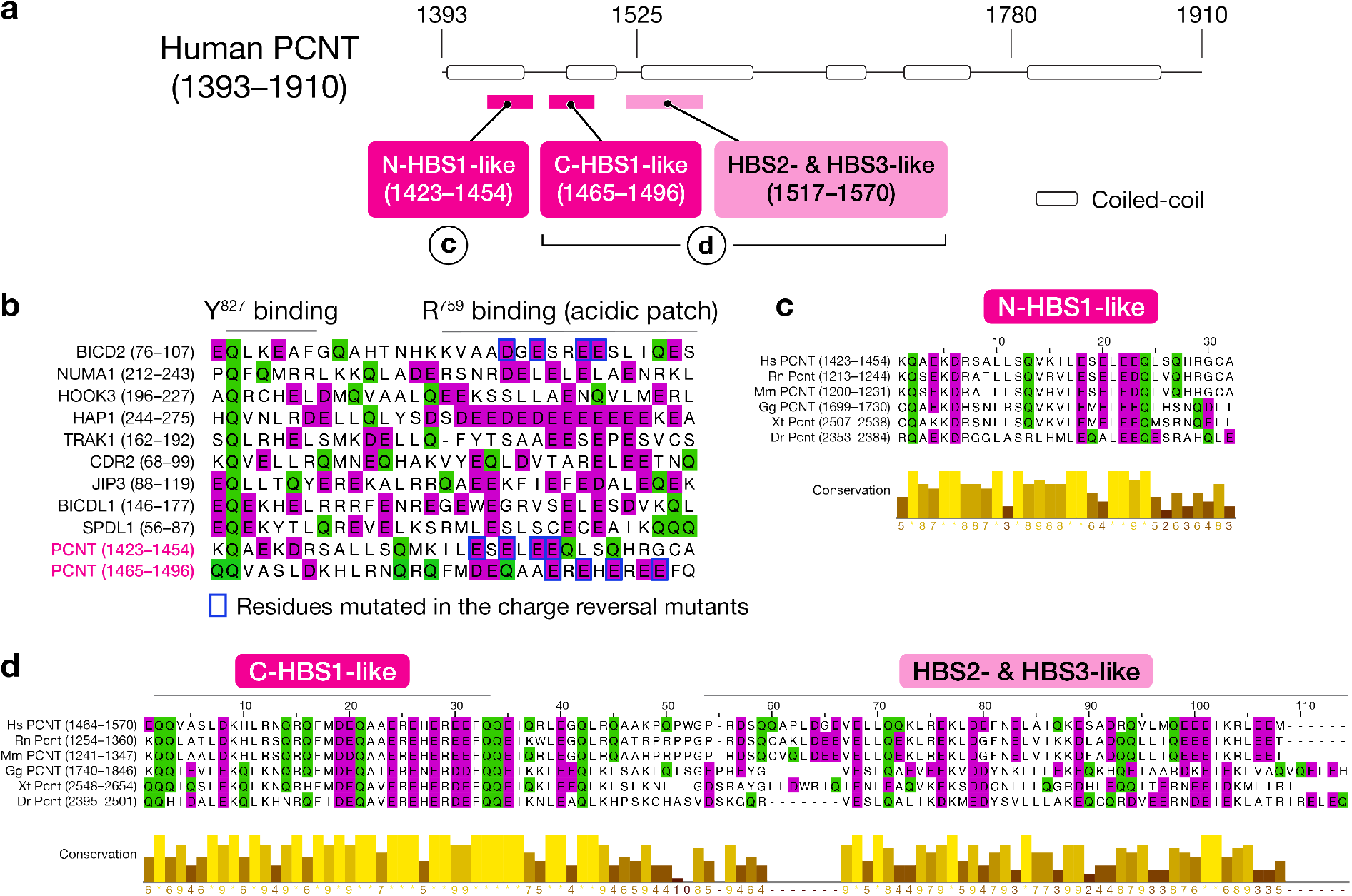
PCNT has HBS1-, HBS2-, and HBS3-like motifs. **a**, Domain structure of human PCNT (1393–1910), showing the HBS1-, HBS2-, and HBS3-like motifs (residue ranges in parentheses). Coiled-coil regions were predicted with Ncoils (Lupas et al., 1991). **b**, Multiple sequence alignment of the HBS1 and HBS1-like motifs of nine dynein cargo adaptors and the two HBS1-like motifs of PCNT (N-HBS1-like and C-HBS1-like). The approximate regions that interact with the dynein heavy chain (DHC) residues Y^827^ and R^759^ are indicated. Blue rectangles mark the residues mutated in the charge-reversal mutants of BICD2N and PCNT (1393–1910) analyzed in the peroxisome transport assay (**Fig. 5**). **c**, Multiple sequence alignment of the N-HBS1-like motif of six vertebrate PCNT orthologous proteins. **d**, Multiple sequence alignment of the C-HBS1-like, HBS2-like, and HBS3-like motifs of the same six PCNT orthologous proteins. Hs, *Homo sapiens*; Rn, *Rattus norvegicus*; Mm, *Mus musculus*; Gg, *Gallus gallus*; Xt, *Xenopus tropicalis*; Dr, *Danio rerio*.

**Extended Data Figure 4.**
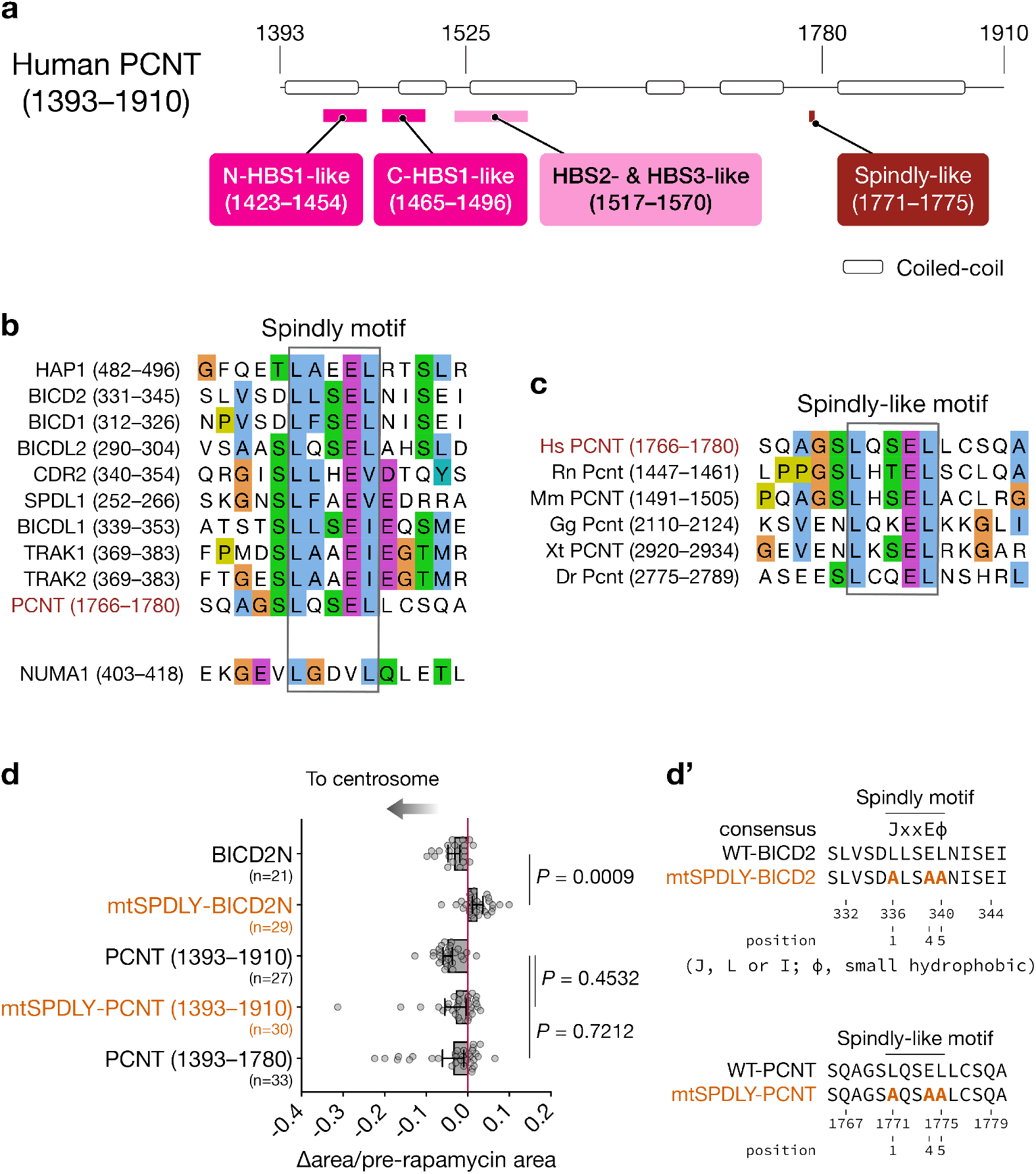
Effects of point mutations in the Spindly and Spindly-like motifs of BICD2 and PCNT (1393–1910) on peroxisome trafficking to the centrosome. **a**, Domain structure of human PCNT (1393–1910), showing the HBS1-, HBS2-, and HBS3-like motifs and one of the putative Spindly-like motifs (residue ranges in parentheses). Coiled-coil regions were predicted with Ncoils (Lupas et al., 1991). **b**, Multiple sequence alignment of the Spindly and Spindly-like motifs of ten dynein cargo adaptors and PCNT. **c**, Multiple sequence alignment of the Spindly-like motif of six vertebrate PCNT orthologous proteins. **d**, Quantification of peroxisome motility. Each data point is the change in mSG1-positive area per cell (15 min post-minus pre-rapamycin) divided by the pre-rapamycin mSG1-positive area. Peroxisomes were scored as trafficking if this ratio was negative and the post-rapamycin area was significantly smaller than the pre-rapamycin area (two-tailed *t*-test). Data are mean ± 95% CI from at least three biological replicates. *P* values were calculated by one-way ANOVA relative to the BICD2N or PCNT (1393–1910) condition. **d’**, Locations of the alanine substitutions at positions 1, 4, and 5 of the Spindly and Spindly-like motifs of BICD2 and PCNT. Hs, *Homo sapiens*; Rn, *Rattus norvegicus*; Mm, *Mus musculus*; Gg, *Gallus gallus*; Xt, *Xenopus tropicalis*; Dr, *Danio rerio*.

**Extended Data Figure 5.**
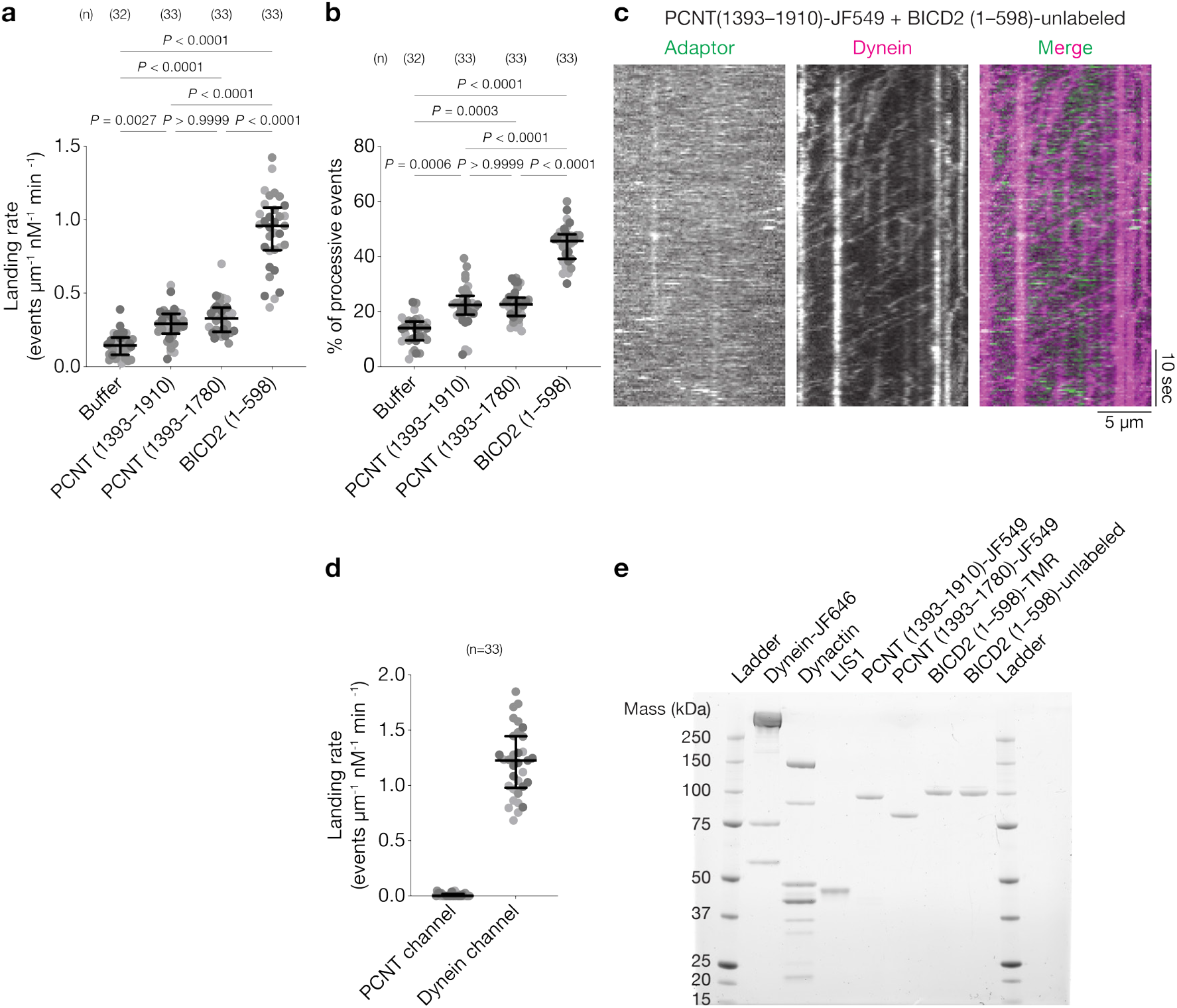
PCNT is a dynein activator (related to Fig. 6). **a**, Landing rate of processive dynein events (events per µm of microtubule per nM dynein per minute). Each spot represents one microtubule, and shades of gray distinguish the three replicates, which came from two independent preparations of adaptor and dynein. Error bars indicate median ± interquartile range. **b**, Percentage of dynein events that were processive. Data are presented as in (**a**). **c**, Representative kymographs of the adaptor, dynein, and merged channels for complexes containing dynein, dynactin, and LIS1 in the presence of PCNT (1393–1910)-JF549 and unlabeled BICD2 (1–598). **d**, Landing rates of processive PCNT (adaptor) and dynein events (events per µm of microtubule per nM dynein per min). Data are presented as in (**a**). **e**, SDS-PAGE of the purified proteins used in the single-molecule motility assays. Statistical significance in **a, b**, and **d** was assessed by a Kruskal-Wallis test with Dunn’s multiple comparisons test. *P* values for pairwise comparisons are shown above the plots, and the number of samples quantified (n) for each condition is shown in parentheses.

## Extended Data Tables

**Extended Data Table 1.** Coiled-coils from UniProt with supporting publications.

| ID | GENE_ID | LENGTH | TYPE | EVIDENCE | POSITION |
| --- | --- | --- | --- | --- | --- |
| D6RGH6 | MCIN_HUMAN | 385 | COILED | ECO:0000269\|PubMed: 24064211 | [178, 227] |
| O00161 | SNP23_HUMAN | 211 | COILED | ECO:0000269\|PubMed: 12556468 | [22, 76] |
| O43715 | TRIA1_HUMAN | 76 | COILED | ECO:0000269\|PubMed: 26071602 | [0, 52] |
| O60239 | 3BP5_HUMAN | 455 | COILED | ECO:0000269\|PubMed: 30217979 | [43, 90] |
| O60239 | 3BP5_HUMAN | 455 | COILED | ECO:0000269\|PubMed: 30217979 | [96, 145] |
| O60239 | 3BP5_HUMAN | 455 | COILED | ECO:0000269\|PubMed: 30217979 | [153, 200] |
| O60239 | 3BP5_HUMAN | 455 | COILED | ECO:0000269\|PubMed: 30217979 | [210, 255] |
| O60271 | JIP4_HUMAN | 1321 | COILED | ECO:0000269\|PubMed: 19644450 | [407, 534] |
| O75030 | MITF_HUMAN | 526 | COILED | ECO:0000305\|PubMed: 24631970 | [354, 402] |
| O75496 | GEMI_HUMAN | 209 | COILED | ECO:0000269\|PubMed: 15260975 | [93, 144] |
| P02671 | FIBA_HUMAN | 866 | COILED | ECO:0000305\|PubMed: 19296670 | [67, 631] |
| P02675 | FIBB_HUMAN | 491 | COILED | ECO:0000305\|PubMed: 10074346 | [156, 222] |
| P08670 | VIME_HUMAN | 466 | COILED | ECO:0000269\|PubMed: 20176112 | [95, 131] |
| P08670 | VIME_HUMAN | 466 | COILED | ECO:0000269\|PubMed: 20176112 | [153, 245] |
| P08670 | VIME_HUMAN | 466 | COILED | ECO:0000269\|PubMed: 20176112 | [302, 407] |
| P12883 | MYH7_HUMAN | 1935 | COILED | ECO:0000255, ECO:0000269\|PubMed: 26150528 | [838, 1935] |
| P23246 | SFPQ_HUMAN | 707 | COILED | ECO:0000255, ECO:0000269\|PubMed: 25765647 | [496, 596] |
| P35658 | NU214_HUMAN | 2090 | COILED | ECO:0000269\|PubMed: 19208808 | [679, 1209] |
| P42224 | STAT1_HUMAN | 750 | COILED | ECO:0000269\|PubMed: 9630226 | [135, 317] |
| P51572 | BAP31_HUMAN | 246 | COILED | ECO:0000269\|PubMed: 23967155 | [164, 237] |
| P51787 | KCNQ1_HUMAN | 676 | COILED | ECO:0000269\|PubMed: 19693805 | [584, 621] |
| Q09013 | DMPK_HUMAN | 629 | COILED | ECO:0000269\|PubMed: 15583383 | [456, 536] |
| Q12933 | TRAF2_HUMAN | 501 | COILED | ECO:0000269\|PubMed: 20385093 | [298, 348] |
| Q13563 | PKD2_HUMAN | 968 | COILED | ECO:0000269\|PubMed: 19556541 | [832, 872] |
| Q13586 | STIM1_HUMAN | 685 | COILED | ECO:0000269\|PubMed: 22451904 | [247, 442] |
| Q13614 | MTMR2_HUMAN | 643 | COILED | ECO:0000269\|PubMed: 16410353 | [592, 627] |
| Q13976 | KGP1_HUMAN | 671 | COILED | ECO:0000269\|PubMed: 16131665 | [1, 59] |
| Q15431 | SYCP1_HUMAN | 976 | COILED | ECO:0000255, ECO:0000269\|PubMed: 29915389 | [738, 798] |
| Q32MZ4 | LRRF1_HUMAN | 808 | COILED | ECO:0000269\|PubMed: 23099021 | [127, 250] |
| Q5HYA8 | MKS3_HUMAN | 995 | COILED | ECO:0000269\|PubMed: 34731008 | [827, 917] |
| Q68DK7 | MSL1_HUMAN | 614 | COILED | ECO:0000269\|PubMed: 23084835 | [212, 282] |
| Q8IZU3 | SYCP3_HUMAN | 236 | COILED | ECO:0000255, ECO:0000269\|PubMed: 24950965 | [65, 223] |
| Q8N4S9 | MALD2_HUMAN | 558 | COILED | ECO:0000269\|PubMed: 28661558 | [438, 548] |
| Q8ND30 | LIPB2_HUMAN | 876 | COILED | ECO:0000269\|PubMed: 21462929 | [100, 313] |
| Q8WWY3 | PRP31_HUMAN | 499 | COILED | ECO:0000269\|PubMed: 17412961 | [84, 120] |
| Q8WWY3 | PRP31_HUMAN | 499 | COILED | ECO:0000269\|PubMed: 17412961 | [180, 215] |
| Q8WYA6 | CTBL1_HUMAN | 563 | COILED | ECO:0000269\|PubMed: 24269683 | [475, 540] |
| Q96D96 | HVCN1_HUMAN | 273 | COILED | ECO:0000269\|PubMed: 20147290 | [222, 266] |
| Q96RU3 | FNBP1_HUMAN | 617 | COILED | ECO:0000269\|PubMed: 17512409 | [66, 259] |
| Q99623 | PHB2_HUMAN | 299 | COILED | ECO:0000269\|PubMed: 28017329 | [189, 238] |
| Q99708 | CTIP_HUMAN | 897 | COILED | ECO:0000269\|PubMed: 34129781 | [34, 84] |
| Q9H9Q4 | NHEJ1_HUMAN | 299 | COILED | ECO:0000269\|PubMed: 18046455 | [127, 170] |
| Q9HC35 | EMAL4_HUMAN | 981 | COILED | ECO:0000269\|PubMed: 25740311 | [13, 63] |
| Q9HCU9 | BRMS1_HUMAN | 246 | COILED | ECO:0000269\|PubMed: 21777593 | [50, 98] |
| Q9HD67 | MYO10_HUMAN | 2058 | COILED | ECO:0000269\|PubMed: 23012428 | [883, 934] |
| Q9P0L9 | PK2L1_HUMAN | 805 | COILED | ECO:0000269\|PubMed: 22193359 | [699, 740] |
| Q9P270 | SLAI2_HUMAN | 581 | COILED | ECO:0000269\|PubMed: 21646404 | [3, 39] |
| Q9UIL1 | SCOC_HUMAN | 159 | COILED | ECO:0000269\|PubMed: 24098481 | [77, 146] |
| Q9ULV4 | COR1C_HUMAN | 474 | COILED | ECO:0000269\|PubMed: 12377779 | [435, 474] |
| Q9UNL4 | ING4_HUMAN | 249 | COILED | ECO:0000269\|PubMed: 22334692 | [24, 118] |

**Extended Data Table 2.**
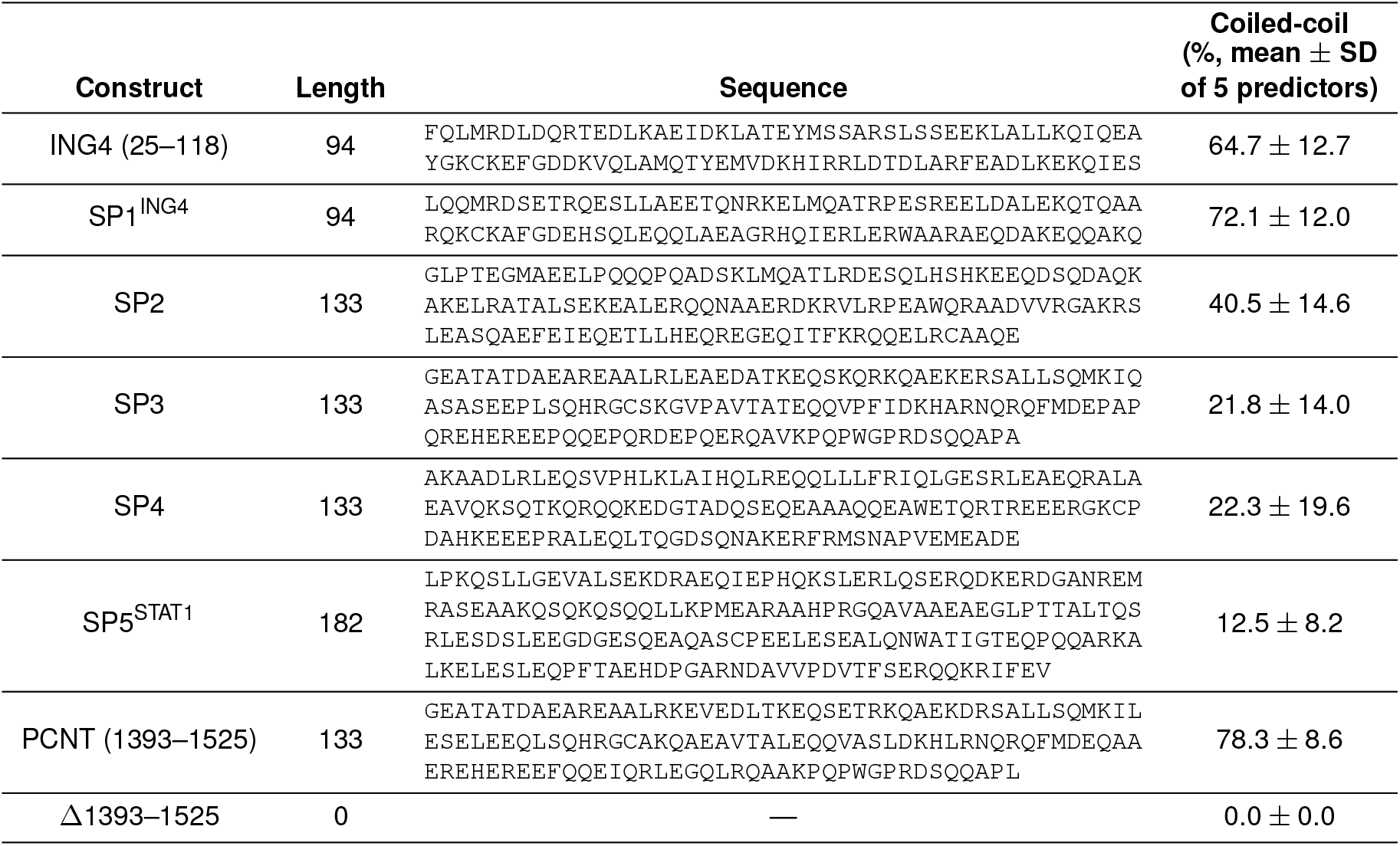
Sequences and predicted coiled-coil contents of PCNT (1393–1525), ING4 (25–118), and the five synthetic sequences. Coiled-coil content is the percentage of residues predicted to be in a coiled-coil, averaged over five predictors (Ncoils, PCOILS, MARCOIL, DeepCoil, and DeepCoil2; settings are given in Methods). *±* indicates the SD across predictors.

**Extended Data Table 3.** Movement of PCNT condensates toward the centrosome after replacement of PCNT (1393–1525). Slopes of linear fits of log_2_ condensate-to-centrosome distance against time for the constructs in **Fig. 3**; more negative slopes indicate faster movement. Movement rate is the slope of each construct divided by the slope of WT, expressed as a percentage. *P* values are from *F*-tests against Δ1393–1525 or WT (— marks a control compared with itself). Rescue was scored from these comparisons: strong rescue, movement significantly faster than Δ1393–1525 and not significantly different from WT; partial rescue, movement significantly faster than Δ1393–1525 but significantly different from WT; and no rescue, movement not significantly faster than Δ1393–1525.

| Construct | Slope | Movement rate<br>(% WT slope) | *P* value<br>vs. Δ1393–1525 | *P* value<br>vs. WT | Rescue? |
| --- | --- | --- | --- | --- | --- |
| Δ1393–1525::ING4 | $-6.919 \times 10^{-4}$ | 16.2 | *P* = 0.0181 | *P* < 0.0001 | No rescue |
| Δ1393–1525::SP1<sup>ING4</sup> | $-3.783 \times 10^{-3}$ | 88.7 | *P* < 0.0001 | *P* = 0.3167 | Strong rescue |
| Δ1393–1525::SP2 | $-2.786 \times 10^{-3}$ | 65.3 | *P* = 0.0015 | *P* = 0.0072 | Partial rescue |
| Δ1393–1525::SP3 | $-3.005 \times 10^{-3}$ | 70.5 | *P* < 0.0001 | *P* = 0.0070 | Partial rescue |
| Δ1393–1525::SP4 | $-1.717 \times 10^{-3}$ | 40.3 | *P* = 0.5533 | *P* < 0.0001 | No rescue |
| Δ1393–1525::SP5<sup>STAT1</sup> | $-1.528 \times 10^{-3}$ | 35.8 | *P* = 0.9863 | *P* < 0.0001 | No rescue |
| WT | $-4.264 \times 10^{-3}$ | 100.0 | *P* < 0.0001 | — | Positive control |
| Δ1393–1525 | $-1.534 \times 10^{-3}$ | 36.0 | — | *P* < 0.0001 | Negative control |

